# Empirical evidence for heterozygote advantage in adapting diploid populations of *Saccharomyces cerevisiae*

**DOI:** 10.1101/033563

**Authors:** Diamantis Sellis, Daniel J. Kvitek, Barbara Dunn, Gavin Sherlock, Dmitri A. Petrov

## Abstract

Adaptation in diploids is predicted to proceed via mutations that are at least partially dominant in fitness. Recently we argued that many adaptive mutations might also be commonly *overdominant* in fitness. Natural (directional) selection acting on overdominant mutations should drive them into the population but then, instead of bringing them to fixation, should maintain them as balanced polymorphisms via heterozygote advantage. If true, this would make adaptive evolution in sexual diploids differ drastically from that of haploids. Unfortunately, the validity of this prediction has not yet been tested experimentally. Here we performed 4 replicate evolutionary experiments with diploid yeast populations (*Saccharomyces cerevisiae*) growing in glucose-limited continuous cultures. We sequenced 24 evolved clones and identified initial adaptive mutations in all four chemostats. The first adaptive mutations in all four chemostats were three CNVs, all of which proved to be overdominant in fitness. The fact that fitness overdominant mutations were always the first step in independent adaptive walks strongly supports the prediction that heterozygote advantage can arise as a common outcome of directional selection in diploids and demonstrates that overdominance of de novo adaptive mutations in diploids is not rare.

---

The most immediate difference between diploids and haploids is that diploids have twice as many gene copies and thus roughly twice as many expected mutations per individual per generation, assuming the same mutation rate per nucleotide. If adaptation is limited by the waiting time for new adaptive mutations, this suggests that diploids might enjoy an adaptive advantage over haploids; indeed, it has been argued that the rate of adaptive evolution in diploids might be up to approximately 1.6 times higher than in haploids (Paquin and Adams 1983; Anderson *et al*. 2004) (although see (Zeyl *et al*. 2003; Gerstein *et al*. 2011)).

Diploids, however, might also suffer an adaptive disadvantage. New mutations in diploids are heterozygous and their effect is thus “diluted” or even completely masked by the presence of the ancestral allele. Unless mutations are fully dominant in fitness, this reduces the probability of fixation of new beneficial mutations and increases the expected frequency that can be reached by deleterious mutations. This fitness effect “dilution” also slows down fixation of adaptive alleles in diploids by roughly two-fold when adaptive mutations are co-dominant in fitness, and by more than that when adaptive mutations are either recessive (they spread in the population slowly) or fully dominant (they suffer a slowdown at high frequencies). This suggests that haploids should gain an advantage in adapting to new environments when the rate of spread of adaptive mutations is the limiting step in adaptation.

These considerations have underpinned a large body of theory (Otto and Gerstein 2008) and generated some specific predictions. One of these is known as Haldane’s sieve and states that adaptive mutations in diploids that do contribute to adaptation are unlikely to be fully recessive in fitness, i.e. adaptive mutations need to be beneficial at least to some extent in heterozygotes (Haldane 1924, 1927; Turner 1981). There is substantial evidence that Haldane’s sieve does operate in evolution, at least to some extent. For instance, fungicide resistance in haploid versus diploid yeast are driven by distinct sets of mutations and at least some of these differences can be attributed to the fact that many of the adaptive mutations in haploids are recessive in resistance and would be invisible in diploids (Anderson *et al*. 2004).

Haldane’s sieve is a claim about the expected fitness of the mutant heterozygote compared to the ancestral homozygote. However, it does not make specific predictions about the fitness of the mutant *homozygote*. Consider a simple Michaelis-Menten model of enzymatic function in which increased catalytic activity is adaptive. A mutation that increases the amount of the enzyme in heterozygotes should generally increase it even further as a homozygote. At first glance this would suggest that such a mutation would thus be even more adaptive as a homozygote and thus partially dominant in fitness. However, consider further that an increase in the level of expression of this enzyme might have additional pleiotropic costs. For example, such costs could be due to elevated levels of protein misfolding, an increased and possibly maladaptive utilization of ribosomes, or additional maladaptive protein-protein interactions. In this case, there might be some intermediate level of expression of the enzyme that balances the costs and benefits in just the right way, with the heterozygote enjoying higher fitness than either ancestral or mutant homozygotes. Such mutations will be overdominant rather than partially dominant in fitness.

How often should we observe beneficial mutations that are not only (partially) dominant but also overdominant in fitness? The answer is currently unknown. We have recently carried out a theoretical study of this question using Fisher’s model of evolution and argued that adaptation should often involve overdominant mutations (Sellis *et al*. 2011). Mutations that are overdominant in fitness are expected to lead to balancing selection and to the persistence of genetic and fitness variation in diploid populations. Intriguingly, our theoretical study suggested that such maintenance of genetic variation in fitness might give diploids an advantage over haploids in changing environments (Sellis *et al*. 2011).

The empirical data on the frequency of fitness overdominance among beneficial mutations in diploids are limited (Hedrick 2012; Guio and González 2015). In a study of fluconazole resistance in yeast, Anderson *et al*. (2004) did not reveal any adaptive mutations that were overdominant in levels of resistance. However, this study focused on fungicide resistance (a component of fitness) rather than on the overall fitness and thus it may have failed to detect fitness overdominance for some of the resistant mutations by overlooking increased pleiotropic costs of the homozygous resistant mutations. Similarly, Gerstein *et al*. (2014) found limited evidence for fitness overdominance in resistance to nystatin in adapting yeast populations. However, these studies potentially miss adaptive mutations unique to diploids by assuming that they acquire beneficial mutations in the same genes as haploids when adapting to the same environmental stress.

Here we perform the first direct test of the adaptive overdominance hypothesis. We experimentally evolved isogenic diploid budding yeast (*Saccharomyces cerevisiae*) populations in glucose-limited continuous cultures (chemostats). Under such conditions, haploid yeast populations have been found to adapt by a small number of adaptive strategies (Kvitek and Sherlock 2011, 2013). One strategy is an approximately 10-fold increase of the copy number of *HXT6* and *HXT7*, which encode high-affinity hexose transporters, providing likely benefit due to the increase of the glucose flux into the cell. An alternative genetic strategy is the disruption of the glucose sensing pathways primarily by the loss of the transcription factor *MTH1*, whose function is to repress the production of high-affinity glucose transporters under the low glucose conditions. The loss of *MTH1* also likely upregulates the expression of *HXT6* and *HXT7* under the low glucose concentrations and thus also likely leads to increased glucose influx. Intriguingly, the combined effect of the loss of *MTH1* and the 10-fold expansion of the *HXT6/7* cluster is deleterious, most likely because the combined effect of both mutations is to increase the amount of hexose transporters beyond the optimal level under these conditions (Kvitek and Sherlock 2011; Chiotti *et al*. 2014).

We find that adaptive mutations in diploids affect the same pathways, however the molecular identity of the mutations is often distinct. Adaptation in diploids is primarily driven by copy number variations (CNVs) in similar genomic regions across evolutionary experiments. We find that the adaptive mutations in diploids are highly beneficial in heterozygotes, but that this benefit is either diminished or even eliminated in homozygotes such that all three adaptive mutations are overdominant in fitness. This result provides to our knowledge the first empirical validation of our adaptive overdominance hypothesis (Sellis *et al*. 2011).

## Materials and Methods

We evolved a diploid yeast strain in replicate glucose-limited continuous cultures (chemostats) until the first adaptive mutations spread in the population (see Results). During our regular plating of populations from the experiments to check for culture contamination, we observed a small colony phenotype when the sample was plated on rich medium (2% glucose YPD). A similar phenotype appeared at different time-points in all chemostats and increased in frequency over time. We sequenced six clones from each chemostat, three of the newly-observed small colony phenotype and three with a wild-type colony phenotype. We then identified the adaptive mutations in each sequenced lineage and proceeded to construct homozygotes for the first adaptive mutations by classical genetics. We measured the competitive fitness of the evolved heterozygotes and homozygotes and ancestral homozygote by competition against a fluorescently labeled reference strain.

### Strains

We evolved a diploid strain of *S. cerevisiae* (GSY2677), which is a derivative of S288C that has an integrated copy of the gene encoding GFP under the *ACT1* promoter integrated at the *YBR209W* gene, achieving constant expression and fluorescence (Kao and Sherlock 2008) (Figure S1). To map our sequenced diploids we used as reference strains GSY1135 and GSY1136 which are haploid clones differing only in the fluorophore they express (DsRed and GFP fluorophores respectively)(Kao and Sherlock 2008). Their genomes have been sequenced and assembled (Kvitek and Sherlock 2011, 2013). We measured competitive fitness against a red ancestral strain derived from the same background. The genealogy of the strains is shown in Figure S1 and all the strains used are listed in Table S1.

### Evolution

Four replicate evolution experiments were carried out in custom-made continuous culture vessels (chemostats) under glucose limitation (0.08% glucose) in Delft medium (Verduyn *et al*. 1992). The total volume was 20ml, the flow rate 0.2 h^−1^ and the temperature was kept constant at 30 °C with a water bath. Daily samples were collected from the outflux of each chemostat. From each sample, 1ml was stored in 25% glycerol at −80 °C, 10 μl were used for cell counting in a Coulter counter, and 10 μl were used for microscope morphological observation and contamination checks. The remaining sample volume (~4ml) was centrifuged at 1,500rpm for 2min on a tabletop centrifuge and the pellet was resuspended in 1ml Sorbitol solution (0.9M sorbitol, 0.1 Tris pH8, 0.1M EDTA pH8) and stored at −20 °C for later DNA preparations. Chemostat samples were also regularly plated on YPD 2% glucose for colony morphology observations and contamination checks. The four chemostats were run continuously without any interruption or change in conditions for 49, 29, 45 and 49 days corresponding to approximately 282,152, 230 and 288 generations respectively.

### Sequencing

Six clones from each chemostat, three clones with the wild-type colony phenotype and three clones with the small colony phenotype on rich medium (2% glucose YPD), were sequenced from samples corresponding approximately to 125 generations of growth (approximately 127, 110, 122 and 123 generations for chemostats 1, 2, 3 and 4 respectively). Samples were bar-coded for multiplexing, pooled and sequenced using the HiSeq 2000 Platform to generate paired end reads of 100nt length. The ancestral clone GSY1136 was also sequenced.

### Mapping reads

All sequence analyses were performed with default parameters unless otherwise noted. We first excluded reads that did not match to the multiplexing barcodes and assigned the rest to their corresponding strain. Reads from each strain were mapped onto the assembled nuclear genome of GSY1135 (Kvitek and Sherlock 2011) using BWA 0.6.1 (Li and Durbin 2010) with trimming parameter −q 10. The mapped reads were then soft-clipped and duplicate Rereads were removed using picard-tools v1.60 (http://picard.sourceforge.net/). Reads were then realigned locally with GATK 1.4–19 (McKenna et al. 2010; DePristo et al. 2011). Putative SNPs and small indels were determined with samtools 0.1.18 (Li et al. 2009) and GATK. The same process was followed for mapping reads to the assembled mitochondrial genome of GSY1135, but with BWA v.0.7.5a, GATK v.2.7–4, picard-tools v.1.101 and samtools 0.1.19.

### Identification of SNPs and small indels

We filtered the putative small indels and SNPs by removing identical variants present in all 4 chemostats and all variants called from regions with coverage less than 30 reads as these most likely represent sequencing or mapping errors. Further filtering was performed by choosing a quality score cutoff for each sequenced strain based on the distribution of quality scores of the particular strain. We modeled the observed distribution of quality scores as a sample from the sum of two random variables, false positives with a Poisson distribution with mean λ, and true positives as a normal distribution with mean *μ* > λ. We used the minimum of the empirical bimodal distributions as a cutoff to further filter the variants. We manually inspected the mapped reads around each variant and narrowed down the true positive SNPs and small indels to a total of 18 across all sequenced lineages based on the quality of the alignments. We removed the remaining false positive polymorphisms by excluding paralogous sequences and polymorphisms shared with the ancestor GSY1136.

Reads from GSY1136 were mapped to the assembled GSY1135 reference and we followed the same procedure as with the evolved strain sequences, but with picard-tools v.1.68 and GATK v.1.6–5. All SNPs and small indels were validated by PCR and Sanger sequencing using the sequencing primers and conditions listed in Tables S2 and S3.

### Identification of large indels and rearrangements

We identified large CNVs (more than 100nt) by visual inspection of changes of the read coverage across chromosomes. We excluded regions with increased coverage that were also present in the ancestral strain and are highly repetitive, such as the rDNA region of chromosome XII (451,450–468,930) and region 212,230–216,200 of chromosome VIII that contains the *CUP1* repeat. We then manually inspected the coverage plots and the orientation and mapping quality of reads at the breakpoints to identify true copy number changes. We estimated the fold change of a CNV by dividing the median read coverage of the CNV region of the focal strain by the median read coverage of the corresponding regions in the sequenced strains from the same chemostat lacking the CNV and normalized by the ratio of the whole genome median coverage of the strains we compared (Table S4). To facilitate the comparison between haploids and diploids we then use twice this value resulting to the fold-change from a haploid genome. For a subset of strains (1.1L, 2.1L, 2.2L, 2.3L, 3.3L and 4.2L) we also performed qPCRs. The rearrangements in chromosome XV aneuploidies were inferred by visual inspection of the mapped reads.

In order to search for possible large scale genomic rearrangements, we used two complementary approaches. First, we implemented a variant of the T-lex algorithm (Fiston-Lavier *et al*. 2011) to search for breakpoints. Briefly, for each strain we calculated the density of orphan reads (read pairs with only one read mapped) across each chromosome. Breakpoints of rearrangements or large indels correspond to regions with increased density of orphan reads with the same orientation. Second, we searched for pairs of reads that map on different chromosomes and compared the results across strains. Putative rearrangements that are shared across strains correspond to erroneous mapping, such as reads that we observed that map on both chromosome VI and chromosome II, which corresponds to the *ACT1* promoter which was used in the integration of GFP. After manually inspecting the putative breakpoints from both methods, no large genomic rearrangements were found.

All nucleotide positions mentioned in the text are relative to the reference strain.

### qPCR

We isolated genomic DNA from clone 2.2L (wild-type colony phenotype from chemostat 2) and from clone I2 (parental clone of chemostat 2) and performed 6 qPCR assays. We repeated the same process but using crude colony lysates instead of genomic DNA and found similar results, thus all the remaining qPCRs were performed using colony lysates. We used primers for the region between *HXT6* and *HXT7* and as a reference gene we used *GLT1* on the same chromosome (Table S5). Quantification was performed in an Eco(TM) Real-Time PCR system running ECO(TM) qPCR Software v0.17.53.0 with iQ SYBR green supermix dye following the protocol and thermal profile listed in Tables S6, and S7. We verified that the primers amplify a single region by PCR and by inspecting the melting curves of the qPCRs. We calculated their efficiency on purified genomic DNA of the ancestral strain in triplicate and across 7 dilutions spanning the ranges used for genotyping. The primers for *GLT1* had efficiency 89.7% (*R^2^ =* 0.935) and the primers for *HXT6* and *HXT7* primers have efficiency 107% (R^2^ = 0.524).

### Resolving the order of mutations

We constructed binary character matrices for each chemostat and included as outgroup the ancestral strain (absence of mutations). We assumed SNPs and indels with identical positions in clones from the same chemostat were identical by descent and coded them as a single character. We also considered as identical by descent CNVs in strains from the same chemostat that span the same exact genes, as breakpoints were in repeated regions and can’t always have a precise location assigned. We performed maximum parsimony analysis with PAUP* v.4.0b10 (Swofford 2003) and parsimony ancestral state reconstruction using Mesquite (Maddison and Maddison 2015). For the ancestral state reconstruction we assumed that mutations observed across all chemostats (*HXT6/7* CNV and chrXV CNV) have a higher rate and thus manually resolve the order of appearance in the clade leading to 1.1L, 1.2L and 1.3L in chemostat 1 and the clade leading to 4.1S, 4.2S, 4.3S in chemostat 4 respectively.

### Genes responsible for phenotypes of clones with CNVs

The CNVs we detected involved a large number of genes. Possibly a subset of them were mostly responsible for the adaptive phenotype and to identify them we searched the *S. cerevisiae* database of GO annotations (Ashburner *et al*. 2000) for genes related to terms ‘detection of glucose’ (GO:0051594) or ‘glucose transport’ (GO:0015758) using SGD (Cherry *et al*. 2012). Additionally, we searched the lists of genes located within CNVs to identify genes found mutated in adaptation to low glucose (Kvitek and Sherlock 2013) or where known from previous studies to be involved in glucose sensing or transport or their regulation.

### Preparation of competition strains

For the construction of strains used in the competitions, we used spores from 4.2L (regular colony phenotype strain from chemostat 4). All spores were mating-type tested and PCR genotyped for the presence or absence of the *UFD2* mutation (see Results); only spores without the mutation were selected. The selected spores were genotyped by qPCR for the presence or absence of the *HXT6/7* CNV and after appropriate matings, homozygous and heterozygous strains for the *HXT6/7* CNV were created (Figure S2). As a reference strain for the competitions, we created a diploid from the same background tagged with DsRed (Bevis and Glick 2002). To do so, we sporulated the ancestral strain GSY2677 and mated the resulting spores with GSY1223, a haploid of the same background tagged with DsRed at the same location (Figure S1). The resulting diploids were sporulated again and the spores were screened under a fluorescence microscope. Spores fluorescing red were mated to construct a diploid homozygous for DsRed (DSY957).

All sporulations were performed by using a sporulation protocol adapted from (Sherman 2002), followed by tetrad dissection using a dissection microscope. Zygotes were picked under a dissection microscope. Successful matings were validated by sporulation and mating-type testing. Mating-type testing was performed by mating with tester strains GSY2476 (*MAT****a***, *met15, ura3, leu2, lys2, his3, xks1*Δ::KanMX) and GSY2670 (*MAT****α****, ura3, leu2, his3, xks1*Δ::KanMX) and then plating on synthetic complete medium with 0.2mg/ml G418 and monosodium glutamate as the nitrogen source.

### Fitness assays

Fitness was determined by replicate competitive fitness assays of homozygote or heterozygote strains for *HXT6/7* CNV in chemostats, using the same conditions and design as the evolutionary experiment. The strains were competed against a diploid reference strain expressing DsRed (DSY957). The strains were first streaked on YPD plates from frozen stock. After four days, the colonies were used to inoculate liquid YPD in culture tubes and incubated overnight in a roller drum at 30 °C. The following morning the chemostats were autoclaved and assembled, and 70% ethanol was run through the pump tubes for 1.5h. Separate starting inocula for each of the fitness competition assays were made by combining 900 μl of DsRed ancestral (DSY957) with 100 μl of each of the GFP strains to be tested, for an initial 9:1 ratio of ancestor to test strain (this allows greater precision in measuring fitness of the test strain). Small volume samples (0.1–0.5ml) were collected one to three times per day from the outflux and stored at 4 °C. Frequencies of competing strains were determined by fluorescence-activated cell sorting (FACS) at the Stanford FACS facility using Scanford, a custom modified FACSScan instrument. A total of 20,000 cells were analyzed per sample. For the GFP-tagged cells we used a 488nm excitation laser and a band pass filter at 525nm (passband 50nm), and for the DsRed-tagged cells an excitation laser at 590nm and a band pass filter at 590nm (passband 20nm). The results were analyzed with FlowJo X 10.0.7r2 (http://www.flowjo.com/), and the frequencies of green and red fluorescing cells was determined. We compared the fluorescence of the samples to controls of green and red cultures of different ages to make sure that differences of fluorescence levels across cell cycle stages or time from sample harvesting did not influence the frequency estimation.

Flow rates were estimated by measuring the volume of water transferred from each pump tube for 1 hour before and after the experiment and averaging the values. The generations were estimated according to (Dykhuizen and Hartl 1983). To estimate selection coefficients, we first calculated the frequency of red and green fluorescing cells and performed a linear regression at the exponential phase of frequency change (Dykhuizen and Hartl 1983).

## Results

We evolved the *S. cerevisiae* diploid strain GSY2677 in four replicate glucose-limited continuous cultures (chemostats) for more than 150 generations (152 to 288). The conditions of growth were chosen to be exactly the same as in the experiments of (Kao and Sherlock 2008) in which populations of a haploid strain (parental to our diploid) were evolved in the same chemostats and using the same media.

Throughout our experiment we regularly measured cell size and population density. We also checked for contamination in part by plating samples from the chemostats on rich medium (2% glucose YPD). We observed that around generation 100 both population density and cell size increased, indicating that adaptive mutations had already spread in the population (Figure S3).

At approximately this same generation time, we began observing the presence of small colonies, seen when plated on rich medium, in the samples from all four chemostats. Note that 50–100 generations was the time required for the first adaptive events in previous evolution experiments using glucose-limited cultures of haploid (Paquin and Adams 1983; Kao and Sherlock 2008) and diploid (Paquin and Adams 1983) populations.

### Mutations in evolved clones

To identify mutations in the evolved clones we performed whole genome sequencing of 24 clones from approximately generation 125: from each of four chemostats we chose three clones with the small-colony phenotype and three with the wild-type colony phenotype.

We mapped the sequenced reads onto a reconstructed ancestral genome, identified mutations, and validated SNPs and small indels by PCR (see Material and Methods for details). On average we found 1.7 mutations per clone. Some mutations were present in multiple clones, both from the same and different chemostats (Table 1). The full list of distinct mutations consists of 10 SNPs, one insertion, and CNVs involving 3 genomic regions (Table 1). The insertion, found in a single clone, is an in-frame single codon addition in the gene *PUF4*. The SNPs include 2 intergenic mutations, 1 synonymous mutation (present in three clones of the same chemostat) and 7 non-synonymous mutations present in 1 to 3 clones (from the same chemostat).

**Table 1.** Mutations identified in the sequenced evolved clones.

| Clone | Gene(s) | Location | Mutation | Amino acid change |
| --- | --- | --- | --- | --- |
| 1.1L | *HXT6/7* | chrIV:1154-1161kb |  | CNV |
|  | *SSN2* | chrIV:1346536 | G → A | Q → stop |
|  | *RPS22A* | chrX:74969 | C → T | D → N |
| 1.2L | *HXT6/7* | chrIV:1154-1161kb |  | CNV |
|  | *SSN2* | chrIV:1346536 | G → A | Q → stop |
|  | *RPS22A* | chrX:74969 | C → T | D → N |
| 1.3L | *HXT6/7* | chrIV:1154-1161kb |  | CNV |
|  | *SSN2* | chrIV:1346536 | G → A | Q → stop |
|  | *RPS22A* | chrX:74969 | C → T | D → N |
| 1.1S | 595 genes | chrXV<sup>a</sup> |  | CNV |
| 1.2S | 595 genes | chrXV<sup>a</sup> |  | CNV |
| 1.3S | 595 genes | chrXV<sup>a</sup> |  | CNV |
| 2.1L | *UTR5* | chrV:85297 | C → A | M → I |
|  | *YFR020W*<sup>f</sup> | chrVI:192542 | C → T | intergenic |
|  | 353 genes | chrIV: dupl: 880-1531kb |  | CNV |
| 2.2L | *HXT6/7* | chrIV:1154-1161kb |  | CNV |
| 2.3L | 353 genes | chrIV: dupl: 880-1531kb |  | CNV |
| 2.1S | 595 genes | chrXV<sup>b</sup> |  | CNV |
| 2.2S | 595 genes | chrXV<sup>b</sup> |  | CNV |
| 2.3S | 595 genes | chrXV<sup>b</sup> |  | CNV |
| 3.1L | *HXT6/7* | chrIV:1154-1161kb |  | CNV |
| 3.2L | *HXT6/7* | chrIV:1154-1161kb |  | CNV |
| 3.3L | *PUF4* | chrVII:467706 | AAT | N (in frame insertion) |
|  | *HXT6/7* | chrIV:1154-1161kb |  | CNV |
| 3.1S | *COG3*<sup>g</sup> | chrV:484675 | G → T | intergenic |
|  | 595 genes | chrXV<sup>c</sup> |  | CNV |
| 3.2S | 595 genes | chrXV<sup>d</sup> |  | CNV |
| 3.3S | 595 genes | chrXV<sup>d</sup> |  | CNV |
| 4.1L | *HXT6/7* | chrIV:1154-1161kb |  | CNV |
| 4.2L | *UFD2* | chrIV:119848 | A → G | V → A |
|  | *HXT6/7* | chrIV:1154-1161kb |  | CNV |
| 4.3L | *HXT6/7* | chrIV:1154-1161kb |  | CNV |
| 4.1S | *CUS1* | chrXIII:750959 | C → T | synonymous |
|  | 595 genes | chrXV<sup>e</sup> |  | CNV |
| 4.2S | *CUS1* | chrXIII:750959 | C → T | synonymous |
|  | 595 genes | chrXV<sup>e</sup> |  | CNV |
| 4.3S | *SNF3* | chrIV:113030 | C → A | S → Y |
|  | *MSH1* | chrVIII:352098 | C → T | T → M |
|  | *CUS1* | chrXIII:750959 | C → T | synonymous |
|  | 595 genes | chrXV<sup>e</sup> |  | CNV |

<sup>a</sup> chrXV CNV chem 1: dupl: 1-323,000; inv: 326,300-326,800; hapl: 326-1,091kb<sup>b</sup> chrXV CNV chem 2: dupl: 1-323,000; inv: 325,200-329,000; hapl: 326-1,091kb<sup>c</sup> chrXV CNV chem 3a: dupl: 1-323,000; hapl: 326-1,091kb<sup>d</sup> chrXV CNV chem 3b: dupl: 1-323,000; hapl: 326-1,091kb<sup>e</sup> chrXV CNV chem 4: dupl: 1-323,000; inv: 325,800-327,350; hapl: 326-1,091kb<sup>f</sup> 200bp upstream<sup>g</sup> 112bp upstream

Heterozygote CNVs were found in all sequenced clones. Based on the fold change (e.g. duplication, haploidization) and the genes they affect we identified three mutation types, two partially overlapping on chromosome IV (which we here refer to as **chrIV CNV** and ***HXT6/7* CNV**) and one on chromosome XV (**chrXV CNV**). The *HXT6/7* CNV evolved at least four independent times, at least once in each chemostat. It is an approximately 10-fold expansion of a ~*7*kb region containing just the *HXT6* and *HXT7* genes, which encode high-affinity glucose transporters (Figure S4). The chrIV CNV, present in two clones from the same chemostat (Figure S5), is a duplication of a large region (~650kb), which includes the *HXT6/7* region that is amplified in the *HXT6/7* CNV. The chrXV CNV also evolved at least four independent times in all four chemostats, and was found in all 12 small-colony phenotype strains (and never in colonies with a wild-type phenotype). It consists of a large partial haploidization of the right arm of chromosome XV and a partial duplication of the left arm of chromosome XV with different breakpoints in each chemostat. With the exception of two strains from chemostat 3, all small-colony strains also had inversions close to the chromosome XV centromere. Although the duplicated and the haploidized regions have distinct breakpoints in each chemostat, all are located within close enough proximity such that the genes affected are exactly the same across all chrXV CNV strains (figure S6, Table 1).

Comparing the mutations we found in the diploid evolutions to mutations in haploid populations of the same strain evolved under exactly the same conditions (Kao and Sherlock 2008; Kvitek and Sherlock 2011, 2013) we find some striking differences. In diploids we found CNVs in all 24 sequenced clones, among which some spanned hundreds of genes. In the haploid strains two of the five sequenced clones had *HXT6* and *HXT7* amplified but no larger CNVs were identified. Also, there is little overlap between genes carrying SNPs in the diploid and haploid evolutionary experiments (Figure S7). For instance, in diploids we didn’t find any loss-of-function mutations in *MTH1*, which was the most frequently mutated gene in the haploid evolutions (sweeping to frequency 23%, 8% and 16% by generation 133 in the three haploid evolutions (Kvitek and Sherlock 2013)), although we observed at least 9 independent likely adaptive events (4 *HXT6/7* CNV, 4 chrXV CNV, and 1 chrIV CNV). The only shared mutations between haploid and diploid evolutions were the expansion covering the genes *HXT6* and *HXT7*, and mutations in *SNF3*, a transmembrane glucose sensor of low glucose concentration (Gancedo 2008).

### The first adaptive mutations are CNVs

The first mutations that sweep to high frequency and/or fix during adaptation are expected to have the largest fitness effect (Orr 1998; Sellis and Longo 2014). Subsequent steps are expected to have smaller fitness effects and compensate for deleterious pleiotropic changes caused by the first mutations (Hindré *et al*. 2012). In the following analysis, we only focus on the first mutations appearing in the sequenced diploid clones. Thus, we are able to unambiguously measure fitness differences and be confident that the adaptations are responses to glucose limitation and were adaptive relative to the ancestral strain in these conditions.

We identified the first adaptive mutation in each lineage by parsimony analysis. First, we constructed a binary character matrix for each chemostat. We assumed that SNPs in clones from the same chemostat that have identical genomic coordinates are identical by descent. We further assumed that CNVs were identical by descent within each chemostat if they involved the duplication and/or deletion of the exactly same genes and their breakpoints were identical, to the precision allowed by our analysis.

We then performed maximum parsimony analysis on the character matrices to reconstruct one tree for each chemostat and to reconstruct the ancestral character states (Figure 1). The first mutation appearing in each strain was unambiguous in all besides two cases, the branch leading to 1.1L, 1.2L and 1.3L in chemostat 1 and the branch leading to 4.1S, 4.2S and 4.3S in chemostat 4. We resolved the order of mutations in these two cases using the observation that across all chemostats *HXT6/7* CNV and chrXV CNV have a higher rate of independent appearance (found in all four chemostats) and are thus likely to have appeared before the SNPs found in the same lineages.

**Figure 1.**
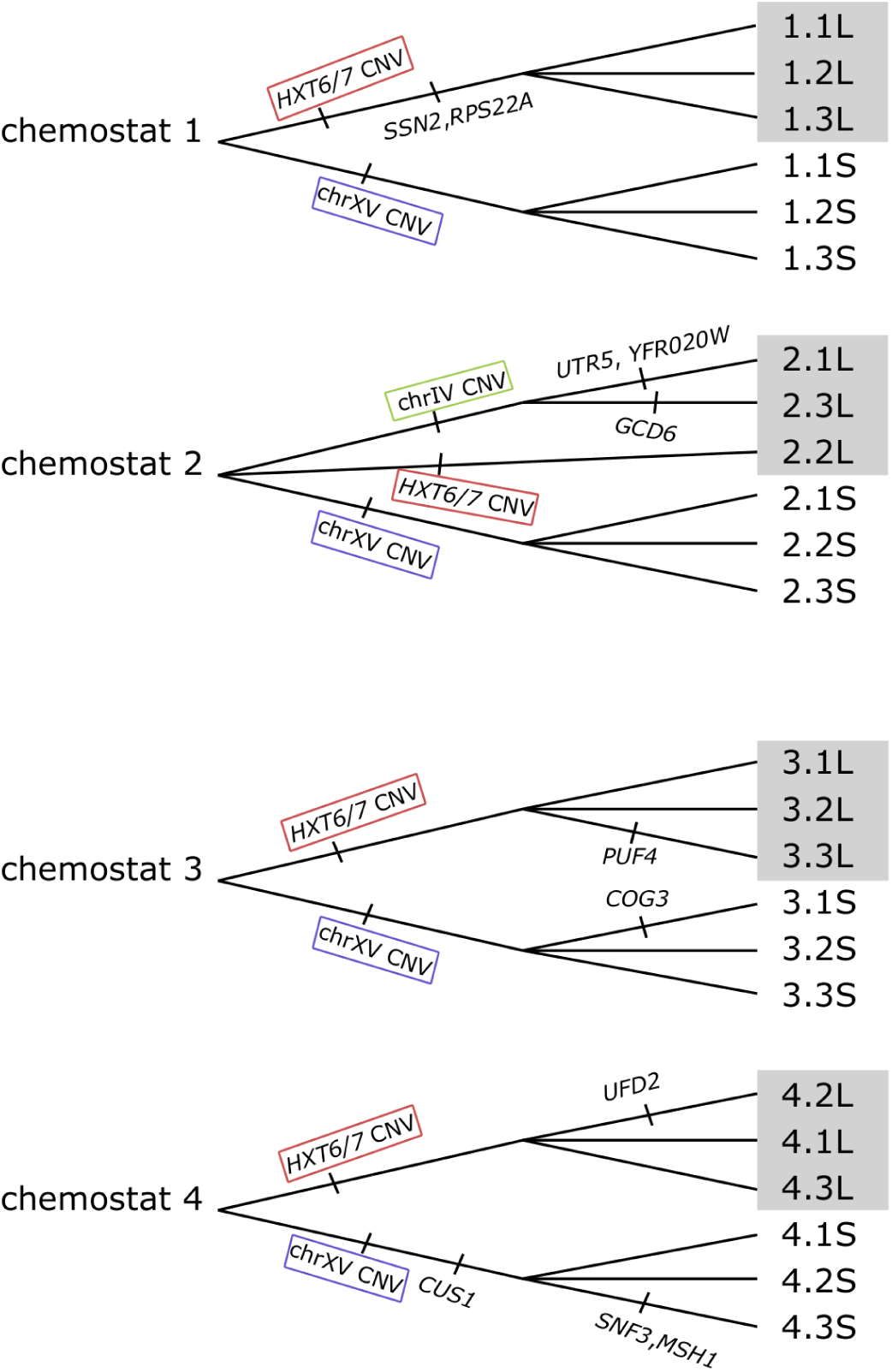
In the sequenced lineages the first adaptive mutations appeared multiple times. The first adaptive mutation in each clone is the expansion of genes *HXT6* and *HXT7* (*HXT6/7* CNV), the partial duplication of chromosome IV (chrIV CNV), and the chromosome XV partial duplication and haploidization (chrXV CNV). The shaded clones have the wild-type colony phenotype.

### Genes affected by CNVs

To understand the mechanisms by which the mutations confer a fitness advantage we explored the genes located in the adaptive CNVs. The chrXV CNV is a complex event involving a large partial deletion and a partial duplication of one copy of chromosome XV. The regions haploidized and duplicated are approximately the same across strains although the boundaries are not identical across chemostats (Table 1, Figure S6). In all cases the haploidized region affects 393 genes, among which are 69 essential genes (Deutschbauer *et al*. 2005), and OSW1 and SSP2, which are necessary for sporulation (Sarkar *et al*. 2002; Li *et al*. 2007). Also found among the haploidized genes is *STD1*, an *MTH1* paralog (Kellis *et al*. 2004) involved in glucose sensing (Gancedo 2008). The duplicated region affects 174 genes (including 25 essential ones) among which is the putative hexose transporter *HXT11*, and *IRA2* which negatively regulates glucose sensing. 300 of the haploidized genes in chrXV CNV and 60 of the duplicated ones were also found haploidized and duplicated respectively in a clone (E7) previously evolved in glucose-limited chemostats (Paquin and Adams 1983; Dunham *et al*. 2002). Also, clones where the distal end of the left arm of chromosome XV was amplified had an increased fitness in glucose-limited chemostats (Sunshine *et al*. 2015).

The chrIV CNV is a transposon mediated partial duplication of chromosome IV including 334 genes, among which are 63 that are essential (Deutschbauer *et al*. 2005), and 3 genes that are deleterious when overexpressed (Tomala and Korona 2013). Also, among the duplicated genes are the negative regulator of glucose transport *MTH1* and the glucose transporters *HXT3, HXT6*, and *HXT7* (Table 2). The same region was previously found expanded in an evolved strain (E8) in glucose-limited chemostats (Paquin and Adams 1983; Dunham *et al*. 2002). Also, among the clones with high fitness in glucose-limited chemostats screened by (Sunshine *et al*. 2015) were those with a partial amplification of the right arm of chromosome IV.

**Table 2.** Properties of the first adaptive mutations.

|  | chrXV CNV | chrIV CNV | *HXT6/7* CNV |
| --- | --- | --- | --- |
| Type | Deletion/Duplication | Duplication | Expansion |
| Genes affected | 393/174 | 334 | 2 |
| Essential genes affected | 69/25 | 63 | 0 |
| Repeatability | 12 clones 4 chemostats | 2 clones 1 chemostat | 10 clones 4 chemostats |
| Genes involved in glucose sensing | *STD1/IRA2* | *MTH1* |  |
| Genes involved in glucose transport |  | *HXT3, HXT6, HXT7* | *HXT6, HXT7* |
| Haploid lethal | Yes | Yes | No |

The *HXT6/7* CNV is an approximately 10-fold expansion of the region including the high-affinity glucose transporters *HXT6* and *HXT7*. These two genes have also been found expanded in previous evolutionary experiments of both haploid (Gresham *et al*. 2008; Kao and Sherlock 2008) and diploid (Brown *et al*. 1998; Gresham *et al*. 2008) strains adapting to glucose-limited continuous cultures.

### Heterozygote advantage

In order to test for heterozygote advantage we focused on the three mutations that first evolved in the sequenced clones: the chrXV CNV, the chrIV CNV and the *HXT6/7* CNV. We expect that these were adaptive responses to glucose limitation and beneficial compared to the ancestral strain.

We attempted to create homozygote mutants for each of the three mutations. Despite our attempts to sporulate the 12 strains carrying the chrXV CNV, none formed spores, suggesting that chrXV CNV is a dominant sterile mutation. Given that 69 essential genes are deleted in the chrXV CNV, it is virtually certain that chrXV CNV is a recessive lethal mutation.

We were successful in producing viable spores from the two lineages carrying the chrIV CNV, but, out of 27 asci dissections, a maximum of only two viable spores per ascus were ever recovered with an average of 1.4 viable spores per dissection, indicating that a recessive lethal mutation was segregating in the spores. We genotyped five surviving spores from three dissections by qPCR and found no instances of the chrIV CNV (If it was not a recessive lethal the probability of the genotyping results would be less than 0.001). We infer that chrIV CNV is lethal as a haploid and likely as a homozygote diploid as well.

Finally, the evolved clones carrying the third mutation tested, the *HXT6/7* CNV, sporulated successfully. We selected four spores of the same evolved strain (4.2L) that had all combinations of mating types and presence-absence of *HXT6/7* CNV but not the mutation in *UFD2*. We performed appropriate matings (Figure S2) to create a homozygote for the *HXT6/7* CNV (*HXT6/7* / *HXT6/7*), a heterozygote (*HXT6/7* / -) and a reference with no *HXT6/7* CNV expansion (-/-). As the evolved strain was tagged with a green fluorescent protein (GFP), we constructed a diploid strain (DSY957) of the same background expressing a red fluorescent protein (DsRed) appropriate for pairwise competitions (Bevis and Glick 2002; Kao and Sherlock 2008). We competed homozygous and heterozygous *HXT6/7* CNV mutants, as well as the reference strain marked with GFP against the DsRed-marked reference strain. The inclusion of the wild-type GFP reference ensured that there was no secondary mutation with significant fitness effect we failed to characterize and allowed us to validate that the fluorophores did not differentially affect fitness. We performed five biological replicates of each set of three pairwise comparisons (red reference vs. green reference, red reference vs. green *HXT6/7* CNV homozygote, red reference vs. green *HXT6/7* CNV heterozygote). We also included one extra control competition of red vs. green reference for a total of 16 competition experiments and measured the frequency of competing strains by fluorescence-activated cell sorting (FACS) (Figure S8, Table S9). In each group of pairwise comparisons we used the same DsRed reference clone in order to minimize non-genetic variations in fitness. The heterozygote *HXT6/7* CNV clone was significantly more fit than both the homozygote *HXT6/7* CNV (p-value = 0.0173, paired t-test) and the ancestral homozygote (p-value = 0.0015). The homozygote *HXT6/7* CNV clone was not significantly different from the ancestral clone in terms of fitness (Figures 2, S9, Table S8).

**Figure 2.**
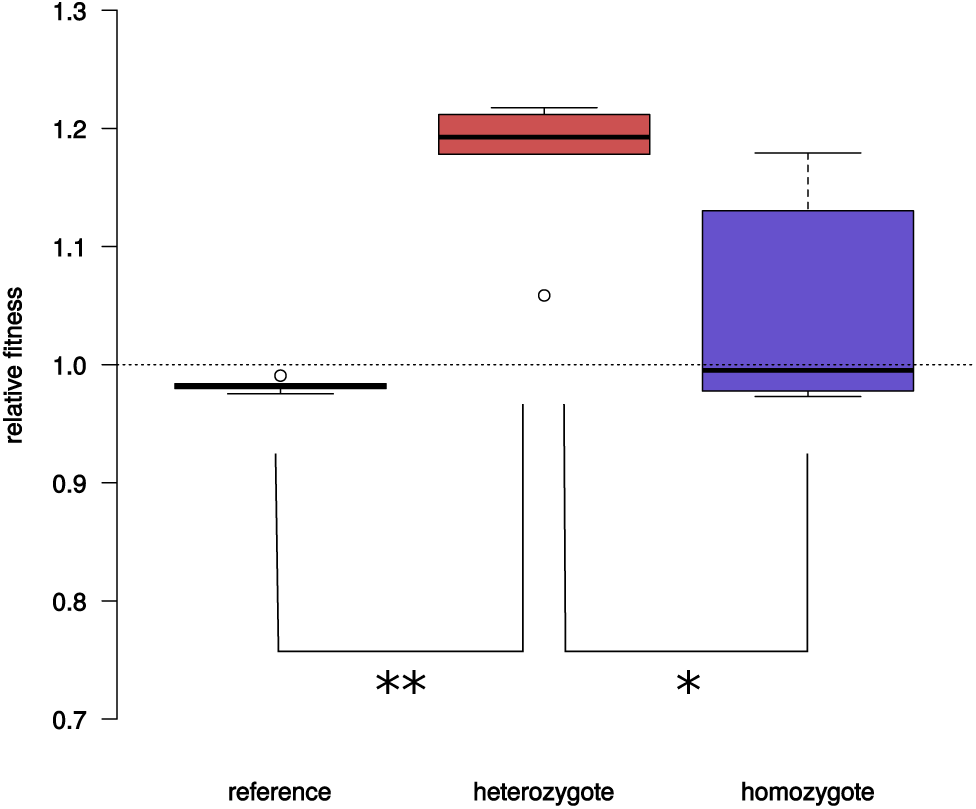
Heterozygote advantage in the expansion of the *HXT6/7* CNV. Relative fitness from pairwise competitions against a common reference strain. The heterozygote *HXT6/7* clone was significantly more fit than both the homozygote *HXT6/7* (p-value = 0.0173, paired t-test) and the ancestral homozygote (p-value = 0.0015).

## Discussion

Our finding that all three tested adaptive mutations that appeared at least nine independent times in four independent evolutions are overdominant in fitness provides strong support for the overdominance hypothesis of Sellis *et al*. (2011). These results show that overdominance is, at the very least, not an exceptionally rare consequence of directional selection, and that directional selection can indeed drive overdominant alleles into the population and lead to maintenance of genetic variation due to heterozygote advantage.

### Diploids and haploids adapt in genetically distinct ways

Because we used a diploid derivate of the haploid strain utilized in previous experiments under identical conditions by Kao and Sherlock (Kao and Sherlock 2008) we were able to compare the outcomes of evolution in haploids and diploids directly. We found that adaptation in diploids generally followed the same phenotypic and molecular strategies as haploids. However, at the genetic level adaptations proceeded in distinct ways. In every case in the diploids the first adaptive mutation was a CNV. This is in contrast to haploids in which only CNVs involving the expansion of *HXT6* and *HXT7* were ever found. At the same time, haploids commonly adapt by the loss-of-function mutations of *MTH1* (discussed below), which were never observed in the diploid evolutions, likely because they are recessive. Overall, it appears that adaptation in diploids is primarily driven by large CNVs that are (partially) dominant in fitness, while most common adaptive mutations in haploids are recessive loss-of-function mutations (Kvitek and Sherlock 2013).

### The first adaptive mutations can be used to test for the overdominance hypothesis

The key purpose of this study was to test the hypothesis that fitness overdominance is common for new adaptive mutations in diploids (Sellis *et al*. 2011). We focused on the three CNVs appearing first in our evolutions and thus certain to be adaptive relative to the ancestor. The mutations were a large deletion and a duplication on chromosome XV (chrXV CNV), a large duplication on chromosome IV (chrIV CNV), and an expansion of the region around the *HXT6* and *HXT7* hexose transporters (*HXT6/7* CNV). It is important to focus on the mutations that spread early during adaptation because subsequent mutations are not necessarily beneficial relative to the ancestral (against which we compete them in order to measure fitness). Such secondary mutations could be adaptive only when present on a specific background (compensatory mutations), or only when competed against another mutant that already reached high frequency in the population or even perhaps only after a change in the chemostat environment (adaptive niche construction). Only by focusing on the three first adaptive CNVs we can be certain that it is appropriate to ascertain the dominance of adaptive mutations by fitness measurements vis-a-vis the ancestor. In addition, these CNVs appeared recurrently across the chemostats, and the same or largely overlapping CNVs were found in diploid clones evolved under glucose limitation in previous experiments (chrIV CNV and chrXV CNV in (Paquin and Adams 1983; Dunham *et al*. 2002), *HXT6/7* CNV in (Brown *et al*. 1998; Gresham *et al*. 2008)), bolstering our confidence that these were indeed adaptive mutations.

### First adaptive mutations are overdominant in fitness

All three mutations showed fitness overdominance. Two of the mutations – chrXV CNV and chrIV CNV – proved to be lethal as haploids and thus likely lethal as homozygote diploids as well. The fact that these CNVs affect hundreds of essential genes makes the inference of the lethality of mutant homozygote a certainty. Thus these two mutations exhibit extreme overdominance, displaying both dominant beneficial and recessive lethal effects. In the third case, that of the *HXT6/7* CNV, the heterozygote is strongly beneficial while the homozygote appears indistinguishable in fitness from the ancestor. Although the number of tested mutations was small, the fact that fitness overdominance was observed in all three tested cases makes it sufficient to claim that fitness overdominance is not an exceedingly rare phenomenon at least for the mutations of large fitness effect.

### Adaptive strategies of the first mutations

While the adaptive mutations in diploids and haploids are qualitatively different, they appear to affect similar pathways. Haploids adapting to low glucose are known to evolve by just a few strategies (Kvitek and Sherlock 2011, 2013) including the disruption of the glucose sensing pathways and the direct increase in the number of hexose transporters in the cell membrane (Figure 3). The identity of adaptive mutations in diploids suggests that the same two strategies are likely followed by diploids as well. The clones with the chrXV CNV possibly follow the glucose sensing disruption strategy by downregulating the glucose sensing pathway as among the haploidized genes is the negative regulator of glucose sensing *STD1* (Figure 3). The strategy of increasing the level of glucose transporters is followed by clones with the *HXT6/7* CNV, which evolved independently in all diploid evolutionary experiments. It is not clear which adaptive strategy is followed by the clones with the chrIV CNV. Among the duplicated genes of chrIV CNV are hexose transporters (*HXT6, HXT7, HXT3*) indicating an increased number of hexose transporters in the cell. However, among the duplicated genes is also the negative regulator of glucose sensing *MTH1*, which would tend to decrease the number of membrane glucose transporters (Figure 3).

**Figure 3.**
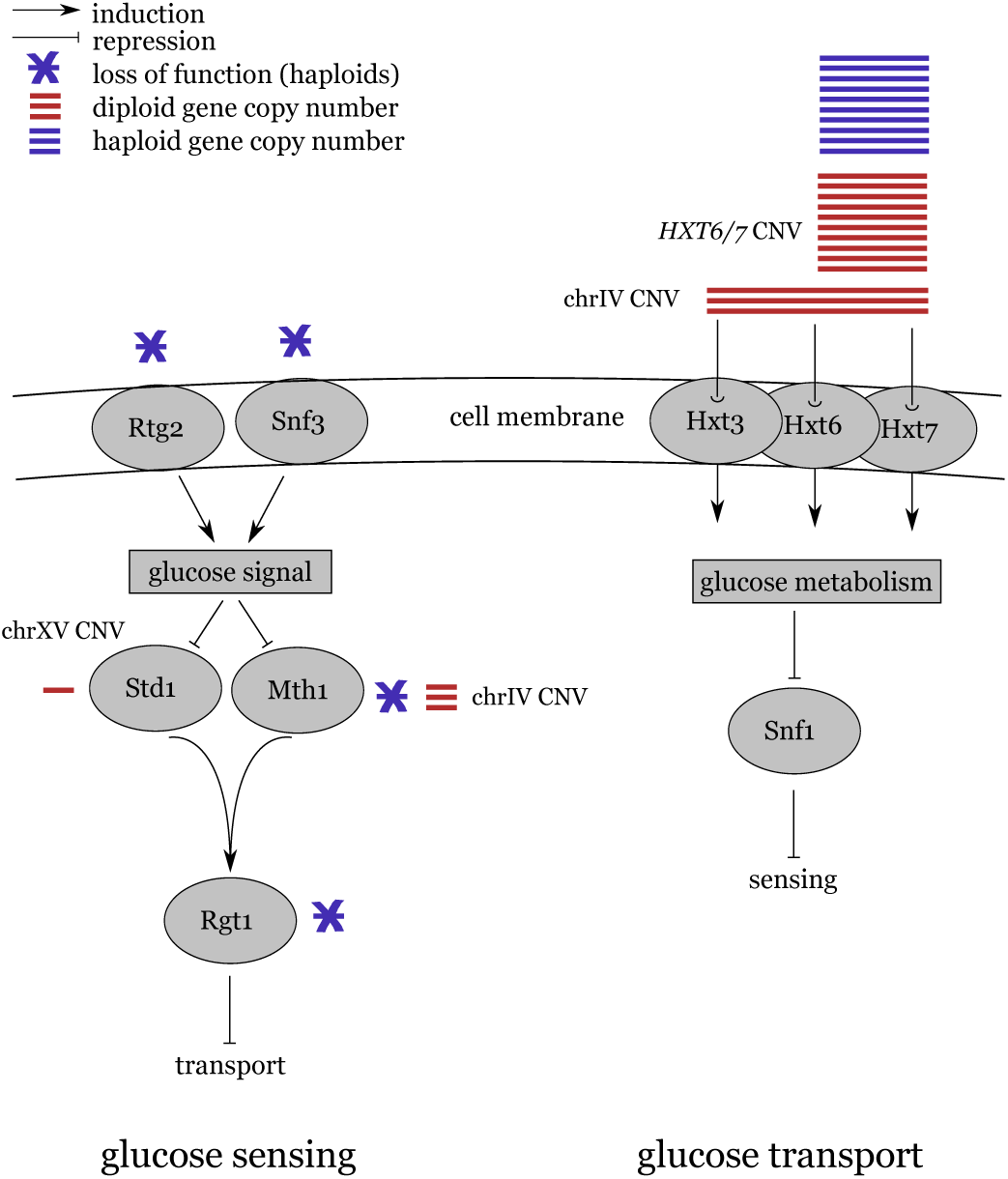
Simplified diagram of glucose sensing and transport pathways annotated to show the major adaptive strategies to growth in glucose-limited environment (adapted from (Kaniak *et al*. 2004)). Adaptation in haploids (blue) include loss of function mutations in the sensing pathway or expansion of the copy number of glucose transporters (Kvitek and Sherlock 2013). Evolved diploid clones (red) had (1) a haploidization of *STD1*, or (2) a duplication of *MTH1, HXT3, HXT6* and *HXT7*, or (3) an expansion of *HXT6* and *HXT7*.

### Epistasis and heterozygote advantage are closely linked

In haploids a negative epistatic relationship was found between loss-of-function mutations in *MTH1* and copy number expansion of the genes *HXT6* and *HXT7* (Kvitek and Sherlock 2011). Each of these adaptive mutations tends to increase the number of hexose transporters in the cell membrane. Although individually adaptive, when combined they generate a strongly maladaptive effect possibly by overshooting the amount of produced Hxt6 and Hxt7 transporters. Consistent with this view, the absolute number of copies of *HXT6* and *HXT7* in the diploid evolved clones with the *HXT6/7* CNV mutation is very close to the copy number found in haploid evolutionary experiments (about 10-fold increase). The close similarity in the fold increase found in haploid and diploid adaptive strains suggests that there may be an optimal level of glucose transporters in the cell membrane shared across haploids and diploids under low glucose conditions, despite the difference in cell size with diploids being larger than haploids (Galitski *et al*. 1999). This is also in agreement with the observation that the relative abundance of Hxt7 in haploids and diploids does not scale as most other proteins (de Godoy *et al*. 2008). It would also explain why we never found both chrXV CNV and *HXT6/7* CNV mutations in the same clone despite finding both in all four chemostats. We thus infer that the existence of an optimum in the number of hexose transporters in both haploids and diploids. This is then likely the cause both of the negative (reciprocal sign) epistasis between *MTH1* and *HXT6/7* duplication in haploids and the overdominance of the *HXT6/7* duplication in diploids.

### Heterozygote advantage is closely related to pleiotropy

As with epistasis, in adaptive mutations there is a close connection of heterozygote advantage and pleiotropy. We initially observed clones with chrXV CNV from their small colony phenotype when plated in rich medium (with high glucose concentration). The mutation is pleiotropic in fitness being adaptive in low glucose but maladaptive in an environment rich in glucose. Such fitness trade-offs are common in yeast (Lang *et al*. 2009; Wenger *et al*. 2011; Sunshine *et al*. 2015). If we consider ploidy as an environment of an allele, as it contributes to the phenotype developed, then our expectation of heterozygote advantage in adapting diploids leads to an expectation of adaptive mutations often being also pleiotropic in the fitness space across environments.

### Large CNVs in adapting diploids

Large CNVs have commonly been found in adapting yeast populations (Adams *et al*. 1992; Dunham *et al*. 2002; Gresham *et al*. 2008; Kvitek and Sherlock 2011; Dunn *et al*. 2012; Hong and Gresham 2014; Payen *et al*. 2014; Sunshine *et al*. 2015). The fitness dominance of large CNVs is the combined fitness effect of multiple linked genes. The overall fitness effect of chrXV CNV and chrIV CNV is possibly the combined effect of multiple deleterious recessive effects that however are masked by dominant beneficial ones. In the homozygotes the benefit from the dominant beneficial mutations is not enough to overcome the effect of the homozygous deleterious mutations and the fitness is decreased. Such a scenario was recently theoretically explored in the case of staggered sweeps caused by linked mutations with zygosity-dependent fitness (Assaf *et al*. 2015). This is a key difference between point mutations and large CNVs that is likely to play out in natural populations as well. Large polymorphic CNVs found in natural populations might be maintained by a similar mechanism of linking dominant beneficial and recessive deleterious mutations by the virtue of being generated by the same CNV such that they cannot be easily unlinked by recombination, as would be the case if they were generated by two independent point mutations.

### Limitations and future directions

Our approach here is limited to only exploring the dominance relationships of the first adaptive mutations as they appeared in our experiment. Partitioning the relative contribution of individual genes among the hundreds in each large CNV is not straightforward. For the chrIV CNV and the duplicated region of chrXV CNV one could possibly measure the fitness of a series of partially overlapping CNVs with the telomeric amplicon approach used by Sunshine *et al*. (2015) and assume that individual gene fitness effects are independent and additive. A complementary approach would be to measure the competitive fitness of single gene knockdowns of the haploidized genes in chrXV CNV (e.g *STD1*).

Our analysis is also limited to only a small number of mutations. We are constrained by the number of direct competitive fitness measurements we can perform and the number of adaptive events we can detect. The detection limit of adaptive mutations can be overcome by an approach based on the barcode technology of Levy *et al*. (2015), which would increase the number of candidate adaptive mutations to hundreds if not thousands.

We do not expect that growth in chemostats under glucose limitation is special in a way that would tend to promote heterozygote advantage. However, we believe it is essential to explore additional environments and species in order to validate the generality of common heterozygote advantage as a consequence of directional adaptation in diploids.

Our results strongly indicate that there is a high probability of fitness overdominance among the first adaptive mutations. As we expect this to have important consequences on the dynamics of adaptation we believe that a comprehensive comparison of adaptation in haploids and diploids is necessary and must involve multiple aspects of adaptation, such as the dynamics, rate of evolution, and size of mutation effects, and must also address the changes of ploidy in life cycles (Orr and Otto 1994; Gerstein and Otto 2009; Gerstein *et al*. 2012; Szövényi *et al*. 2013).

## Acknowledgments

We would like to thank Sophia Christel for helping with making DNA sequencing libraries. Hunter Fraser, Ashby Morrison, Russell Fernald, Peter Zee and Yiqi Zhou provided lab equipment and useful feedback. We would also like to thank Katja Schwartz for advice and guidance and Sandeep Venkataram for invaluable help with setting up the yeast section of the Petrov lab. We also thank Yuan Zhu, Natalia Chousou-Polydouri and the members of the Fraser lab and Petrov lab for feedback. Funding for this project was from a Stanford Graduate Fellowship (DS), a Stanford Center for Evolutionary and Human Genomics (CEHG) Fellowship (DS), and RO1 HG003328 (GS).

---

**Table S1.** List of strains used

| Name | Identifier | Description |
| --- | --- | --- |
| GSY2677 | GSY2677 | Parental genotype |
| I1 | DSY486 | Inoculum of chemostat 1 (GSY2677) |
| I2 | DSY487 | Inoculum of chemostat 2 (GSY2677) |
| I3 | DSY488 | Inoculum of chemostat 3 (GSY2677) |
| I4 | DSY489 | Inoculum of chemostat 4 (GSY2677) |
| red ancestral | DSY957 | DsRed tagged, parental genotype |
| -/- | DSY751 | Parental genotype |
| *HXT6/7* /- | DSY756 | Heterozygote for *HXT6/7* CNV |
| *HXT6/7* / *HXT6/7* | DSY787 | Homozygote for *HXT6/7* CNV |
| 1.1L | DSY436 | Wild-type phenotype, Chemostat 1 |
| 1.2L | DSY437 | Wild-type phenotype, Chemostat 1 |
| 1.3L | DSY438 | Wild-type phenotype, Chemostat 1 |
| 1.1S | DSY439 | Small-colony phenotype, Chemostat 1 |
| 1.2S | DSY440 | Small-colony phenotype, Chemostat 1 |
| 1.3S | DSY441 | Small-colony phenotype, Chemostat 1 |
| 2.1L | DSY442 | Wild-type phenotype, Chemostat 2 |
| 2.2L | DSY443 | Wild-type phenotype, Chemostat 2 |
| 2.3L | DSY444 | Wild-type phenotype, Chemostat 2 |
| 2.1S | DSY445 | Small-colony phenotype, Chemostat 2 |
| 2.2S | DSY446 | Small-colony phenotype, Chemostat 2 |
| 2.3S | DSY447 | Small-colony phenotype, Chemostat 2 |
| 3.1L | DSY448 | Wild-type phenotype, Chemostat 3 |
| 3.2L | DSY449 | Wild-type phenotype, Chemostat 3 |
| 3.3L | DSY450 | Wild-type phenotype, Chemostat 3 |
| 3.1S | DSY451 | Small-colony phenotype, Chemostat 3 |
| 3.2S | DSY452 | Small-colony phenotype, Chemostat 3 |
| 3.3S | DSY453 | Small-colony phenotype, Chemostat 3 |
| 4.1L | DSY454 | Wild-type phenotype, Chemostat 4 |
| 4.2L | DSY455 | Wild-type phenotype, Chemostat 4 |
| 4.3L | DSY456 | Wild-type phenotype, Chemostat 4 |
| 4.1S | DSY457 | Small-colony phenotype, Chemostat 4 |
| 4.2S | DSY458 | Small-colony phenotype, Chemostat 4 |
| 4.3S | DSY459 | Small-colony phenotype, Chemostat 4 |

**Table S2.**
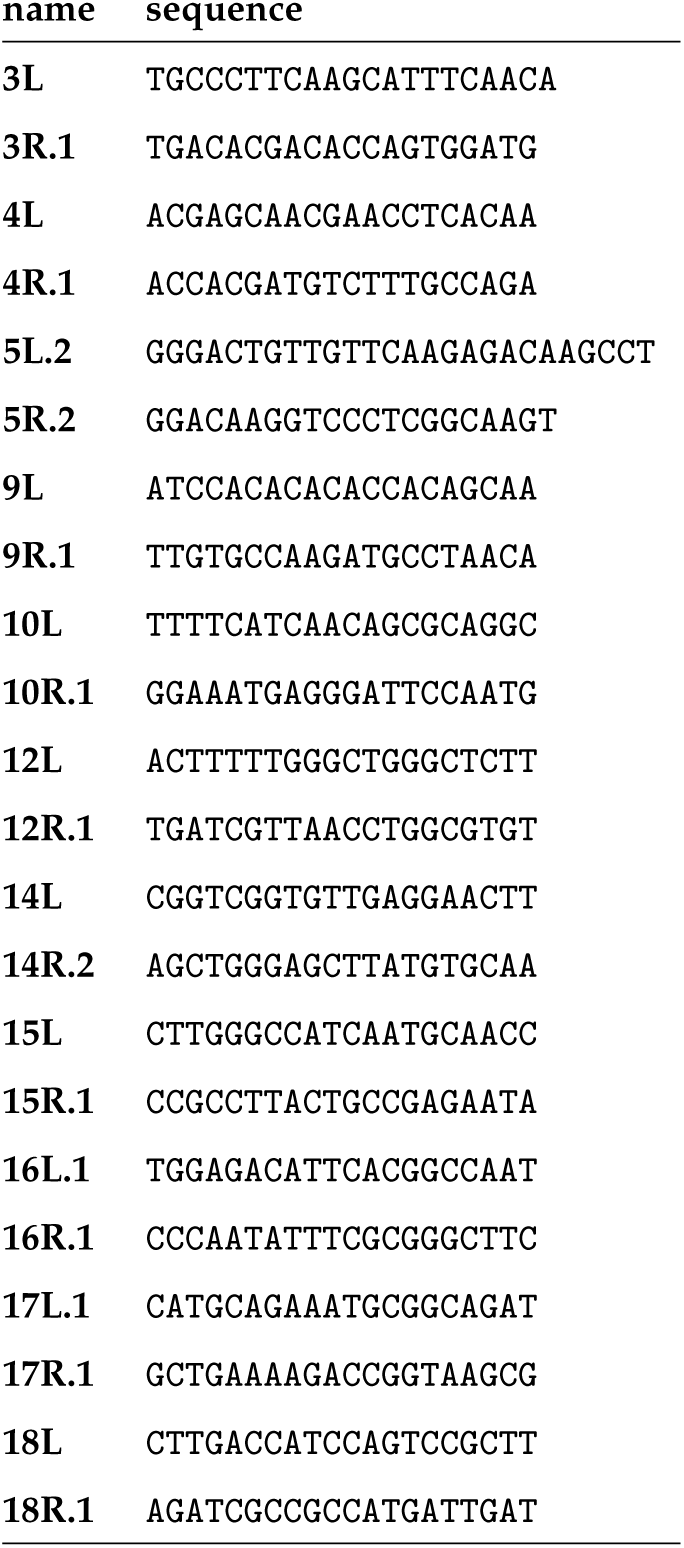
primers used for SNP validation

**Table S3.**
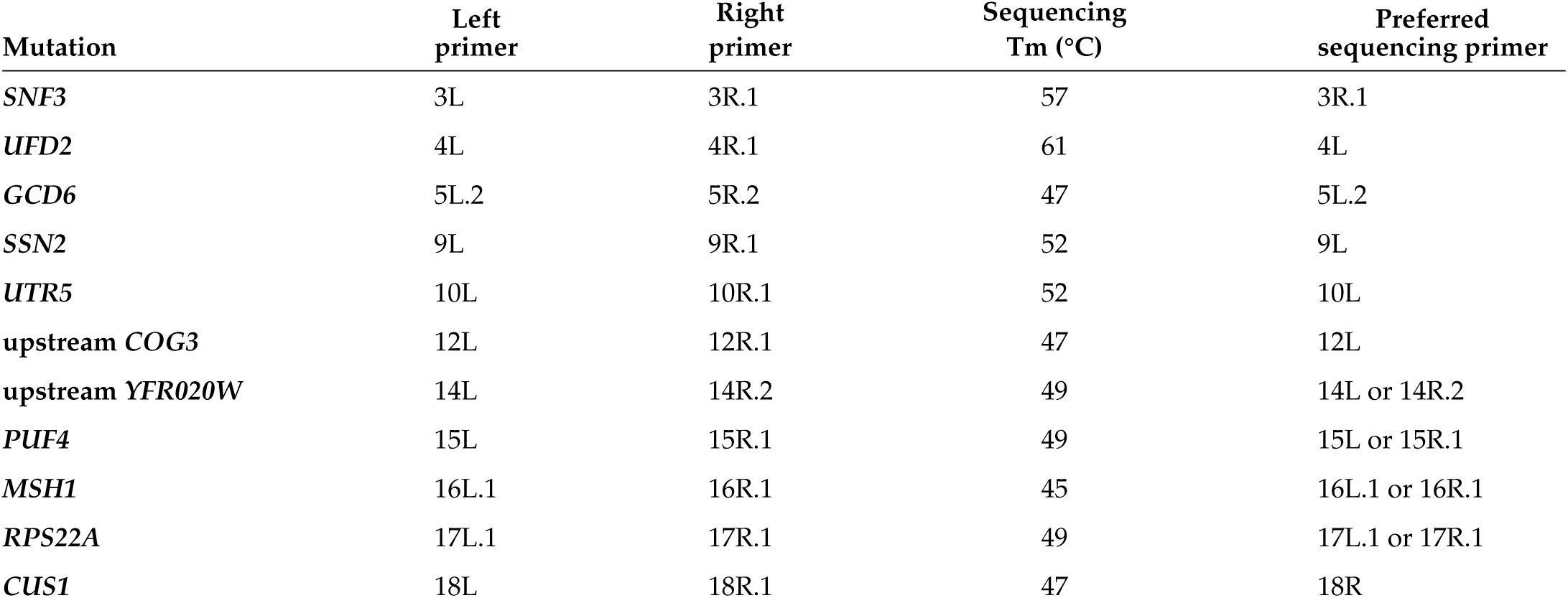
Conditions and preferred primer for PCR

**Table S4.** Relative fold-change of CNVs with respect to the ancestral diploid. We calculated separately the values for chrXV CNV in the duplicated and haploidized region (listed as chrXVd and chrXVh respectively).

| Strain | CNV or CNV Region | Fold-change (mean ± stdev) |
| --- | --- | --- |
| 1.1L | *HXT6/7* | 8.25 ± 2.68 |
| 1.2L | *HXT6/7* | 8.71 ± 2.89 |
| 1.3L | *HXT6/7* | 9.71 ± 3.46 |
| 1.1S | chrXVh | 1.23 ± 0.32 |
| | chrXVd | 3.32 ± 0.61 |
| 1.2S | chrXVh | 1.27 ± 0.34 |
| | chrXVd | 3.40 ± 0.63 |
| 1.3S | chrXVh | 1.13 ± 0.33 |
| | chrXVd | 3.30 ± 0.66 |
| 2.1L | chrIV | 2.60 ± 0.57 |
| 2.2L | *HXT6/7* | 8.94 ± 3.64 |
| 2.3L | chrIV | 2.66 ± 0.54 |
| 2.1S | chrXVh | 1.12 ± 0.30 |
| | chrXVd | 3.36 ± 0.60 |
| 2.2S | chrXVh | 1.31 ± 0.33 |
| | chrXVd | 3.45 ± 0.69 |
| 2.3S | chrXVh | 1.44 ± 0.34 |
| | chrXVd | 3.65 ± 0.69 |
| 3.1L | *HXT6/7* | 9.32 ± 3.10 |
| 3.2L | *HXT6/7* | 8.54 ± 2.51 |
| 3.3L | *HXT6/7* | 13.18 ± 4.33 |
| 3.1S | chrXVh | 1.28 ± 0.41 |
| | chrXVd | 3.37 ± 0.82 |
| 3.2S | chrXVh | 1.56 ± 0.42 |
| | chrXVd | 3.35 ± 0.75 |
| 3.3S | chrXVh | 1.37 ± 0.34 |
| | chrXVd | 3.27 ± 0.70 |
| 4.1L | *HXT6/7* | 8.58 ± 2.83 |
| 4.2L | *HXT6/7* | 8.97 ± 2.60 |
| 4.3L | *HXT6/7* | 8.78 ± 2.40 |
| 4.1S | chrXVh | 1.21 ± 0.37 |
| | chrXVd | 3.27 ± 0.78 |
| 4.2S | chrXVh | 1.45 ± 0.34 |
| | chrXVd | 3.48 ± 0.74 |
| 4.3S | chrXVh | 1.25 ± 0.34 |
| | chrXVd | 3.26 ± 0.69 |

**Table S5.**
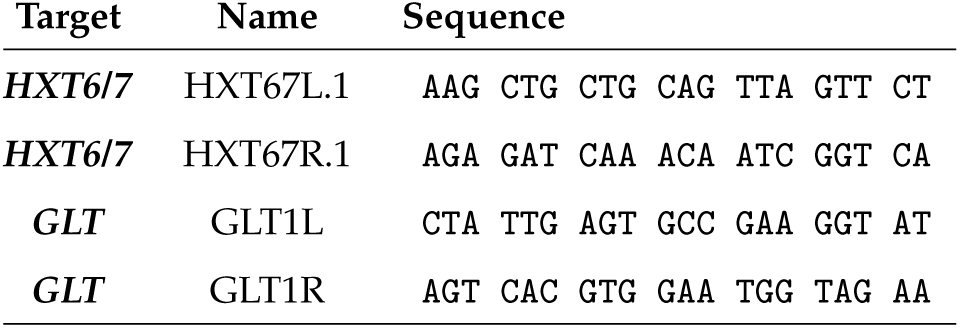
qPCR primers

**Table S6.** qPCR protocol

| Component | Volume (μl) |
| --- | --- |
| iQ mix | 5 |
| $H_2O$ | 3 |
| primer1 10μM | 0.5 |
| primer2 10μM | 0.5 |
| lysate | 1 |
| total | 10 |

**Table S7.** qPCR thermal profile

| Stage | Temperature ($^{\circ}$C) | Duration | Cycles |
| --- | --- | --- | --- |
| polymerase activation | 95 | 00:03:00 | 1 |
| PCR Cycling | 95 | 00:00:15 | 40 |
| PCR Cycling | 57 | 00:00:30 | 40 |
| PCR Cycling | 72 | 00:00:30 | 40 |
| Melt Curve | 95 | 00:00:15 | 1 |
| Melt Curve | 55 | 00:00:15 | 1 |
| Melt Curve | 95 | 00:00:15 | 1 |
| Total Program Length |  | 1:29:00 |  |

**Table S8.** Relative selection coefficients (*s*)

| Biological replicate | Ancestral | Heterozygote | Homozygote |
| --- | --- | --- | --- |
| 1 | 0.9753 | 1.2176 | 1.1302 |
| 2 | 0.9839 | 1.2117 | 0.9951 |
| 3 | 0.9797 | 1.1781 | 0.9777 |
| 4 | 0.9818 | 1.0586 | 0.9730 |
| 5 | 0.9907 | 1.1925 | 1.1792 |

**Table S9.** Experimental design of competitive chemostats

| Biological replicate | Chemostat | Clone 1 | Clone 2 | Competition |
| --- | --- | --- | --- | --- |
| 1 | 1 | DSY957 | DSY751 | red vs. green ancestral |
| 1 | 5 | DSY957 | DSY757 | red vs. green heterozygote |
| 1 | 9 | DSY957 | DSY787 | red vs. green homozygote |
| 2 | 2 | DSY957 | DSY751 | red vs. green ancestral |
| 2 | 6 | DSY957 | DSY757 | red vs. green heterozygote |
| 2 | 10 | DSY957 | DSY787 | red vs. green homozygote |
| 3 | 3 | DSY957 | DSY751 | red vs. green ancestral |
| 3 | 7 | DSY957 | DSY757 | red vs. green heterozygote |
| 3 | 11 | DSY957 | DSY787 | red vs. green homozygote |
| 4 | 4 | DSY957 | DSY751 | red vs. green ancestral |
| 4 | 8 | DSY957 | DSY757 | red vs. green heterozygote |
| 4 | 12 | DSY957 | DSY787 | red vs. green homozygote |
| 5 | 13 | DSY957 | DSY751 | red vs. green ancestral |
| 5 | 14 | DSY957 | DSY757 | red vs. green heterozygote |
| 5 | 15 | DSY957 | DSY787 | red vs. green homozygote |
| 6 | 16 | DSY957 | DSY751 | red vs. green ancestral |

**Figure S1.**
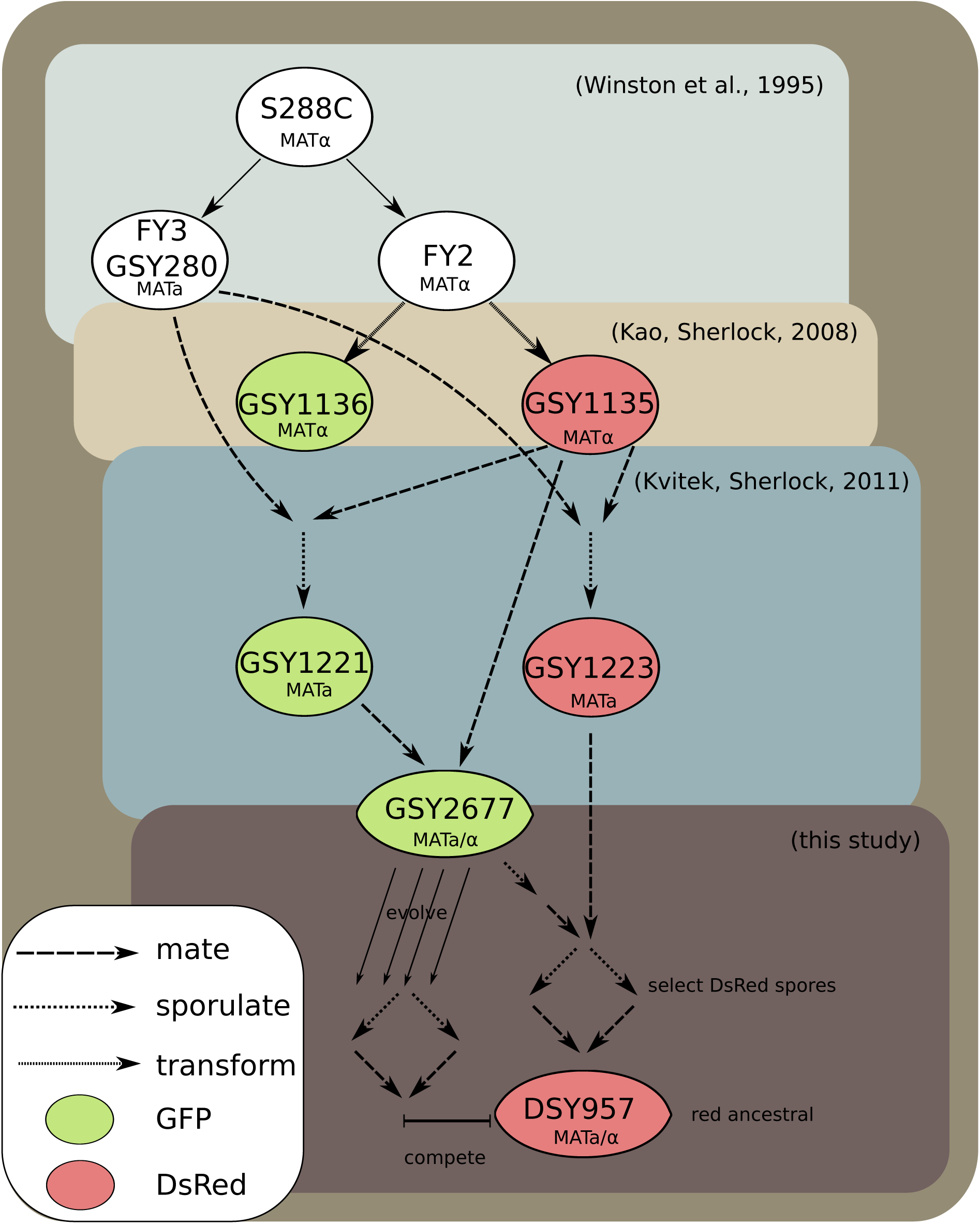
The diploid strain we evolved (GSY2677) is derived from strains used in evolutionary experiments of haploids under the same conditions (Kao, Sherlock 2008) derived from S288C (Winston et al 1995). The red ancestral (DSY957) we used for the competitions is also derived from the same background. We mapped all sequenced strains in the current study against the two previously sequenced genomes of GSY1136 and GSY1135 which are ancestral to the evolved strain.

**Figure S2.**
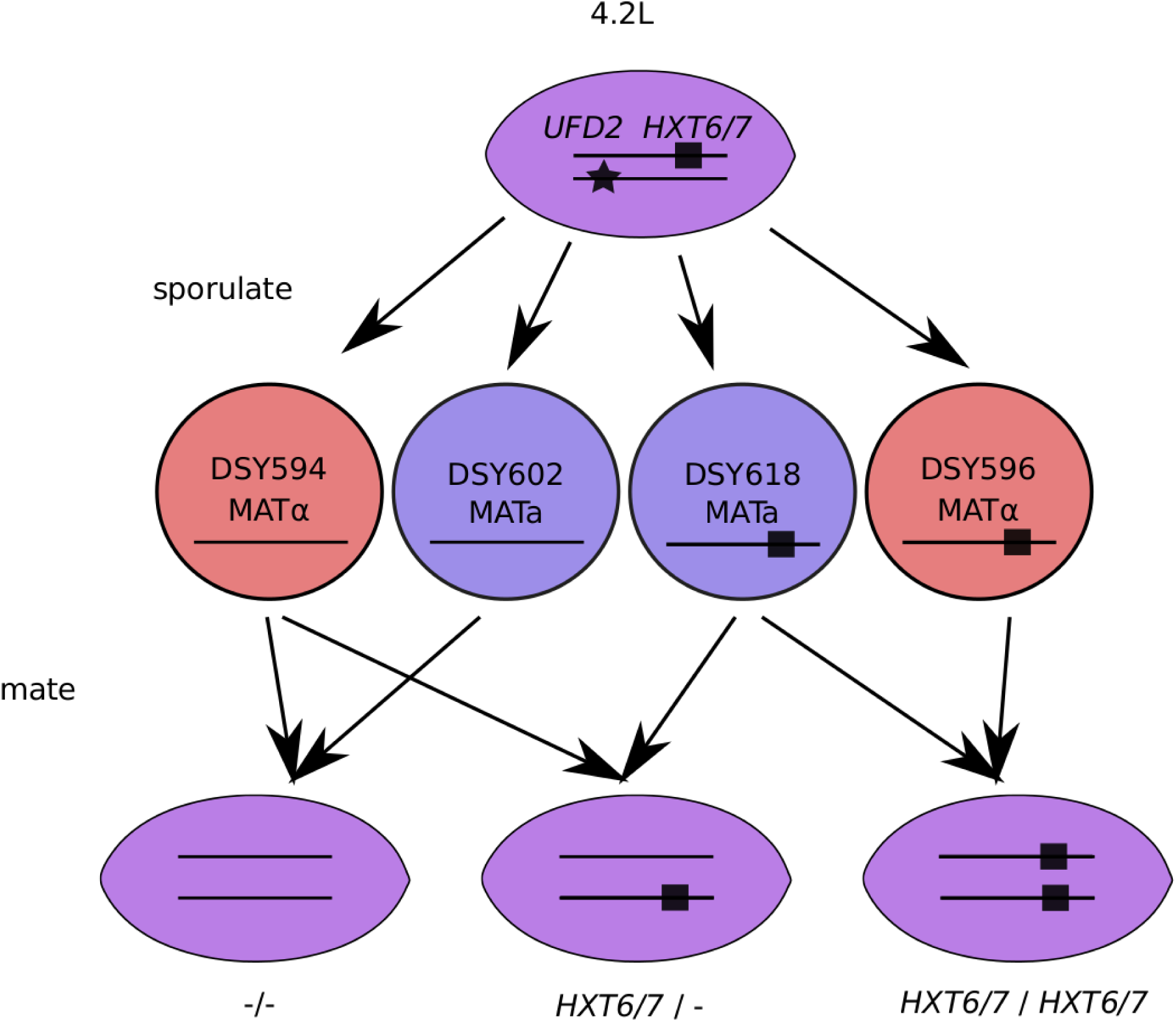
Mating scheme for preparing strains that differ only at a single locus. Evolved strains with the *HXT6/7* CNV were sporulated to create a large number of haploid spores. The spores were genotyped for the *HXT6/7* CNV by qPCR and for secondary mutations, if present in the evolved diploid sporulated, by PCR. We chose four spores from the same evolved strain (4.2L) that had all combinations of mating types and presence-absence of *HXT6/7* CNV. We then performed the appropriate matings and created an ancestral genotype (-/-), a heterozygote for the *HXT6/7* CNV (*HXT6/7* / -) and a homozygote (*HXT6/7* / *HXT6/7*).

**Figure S3.**
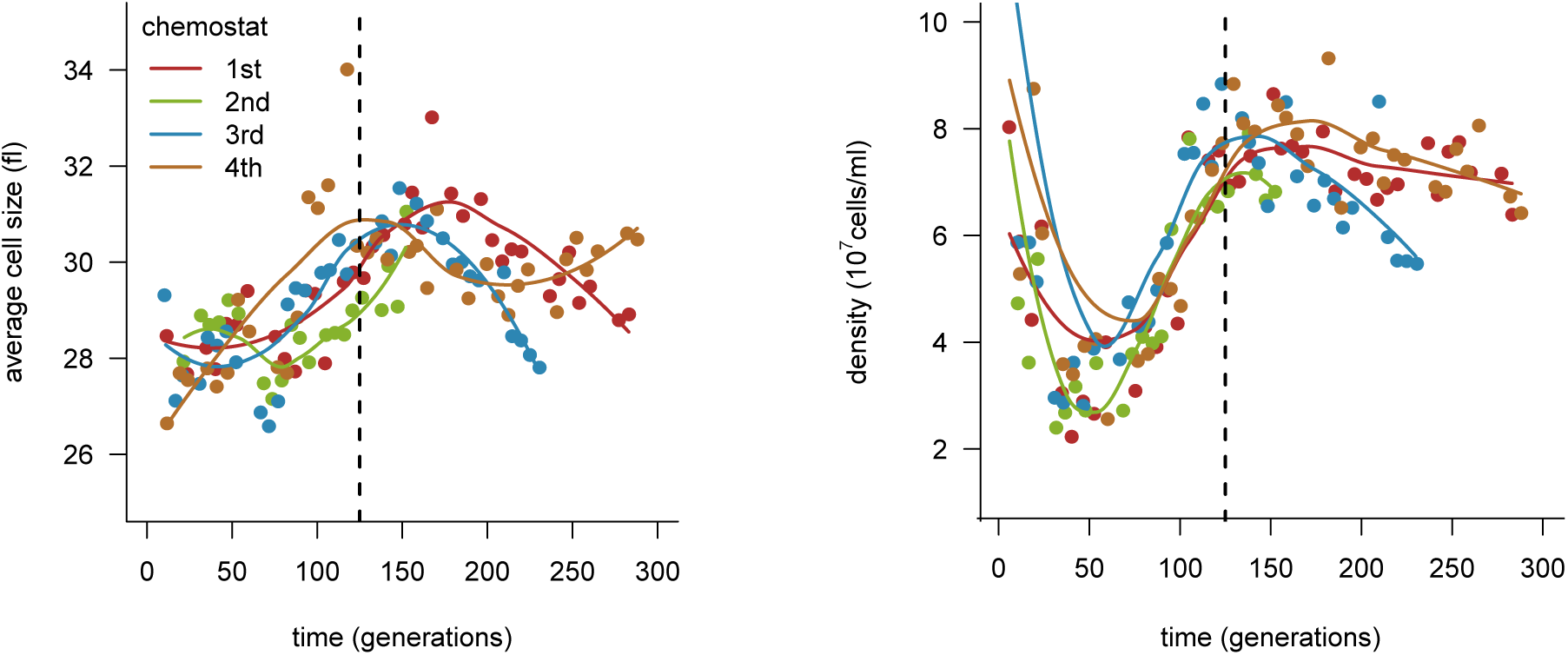
Average cell size and population density during chemostat evolutionary experiments. Each dot corresponds to a unique sample color-coded by chemostat. The dashed vertical line corresponds to generation 125 from which the sequenced clones were sampled. The colored curves are local polynomial fitted lines to help visualize the general trends. At approximately generation 100 average cell size and density is changing in all chemostats. The large initial values of cell density correspond to the equilibration period.

**Figure S4.**
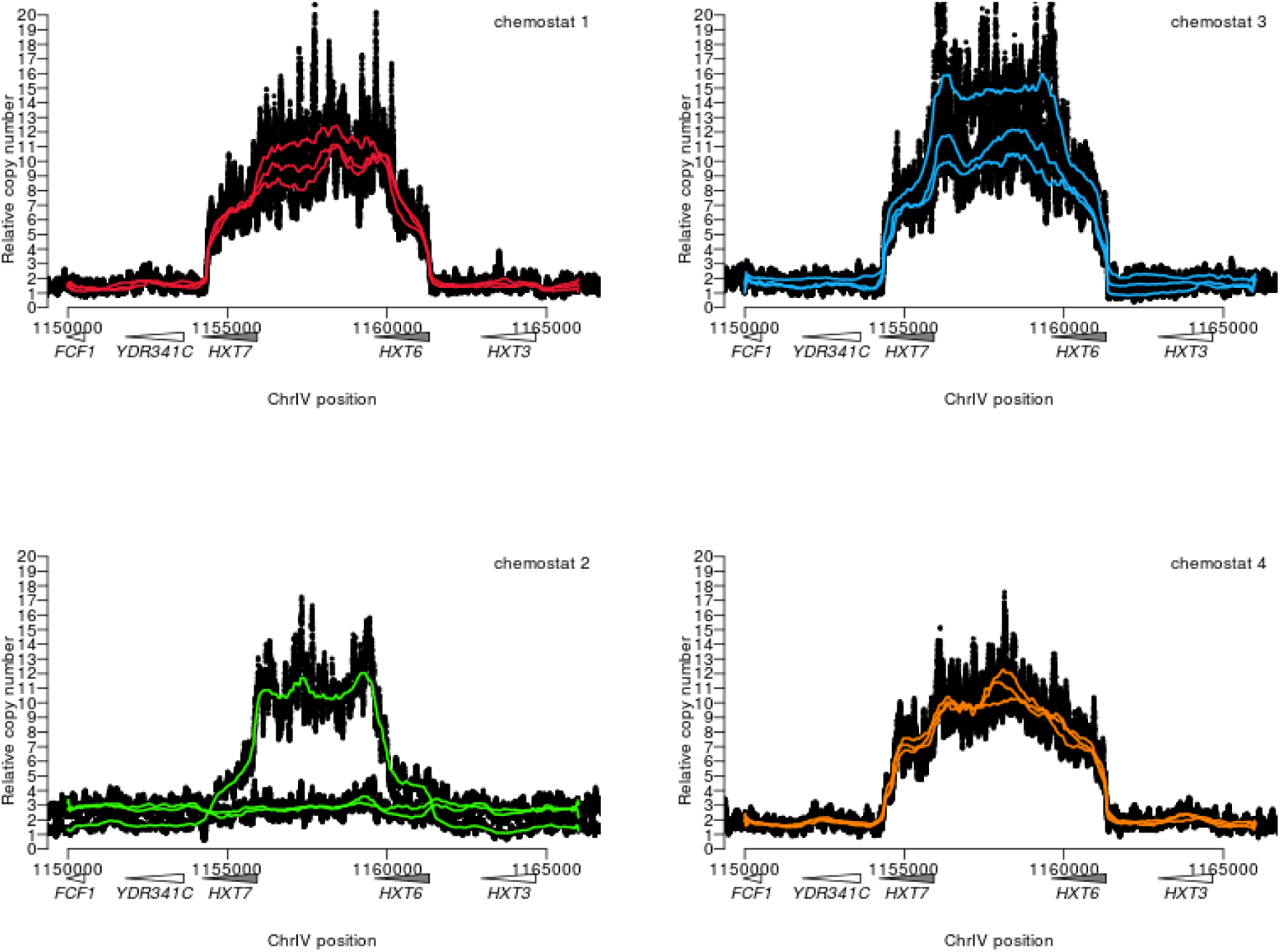
Relative coverage across *HXT6/7* CNV in wild-type colony phenotype strains from each chemostat. Dots correspond to the relative coverage, while continuous lines to running medians of 801bp windows.

**Figure S5.**
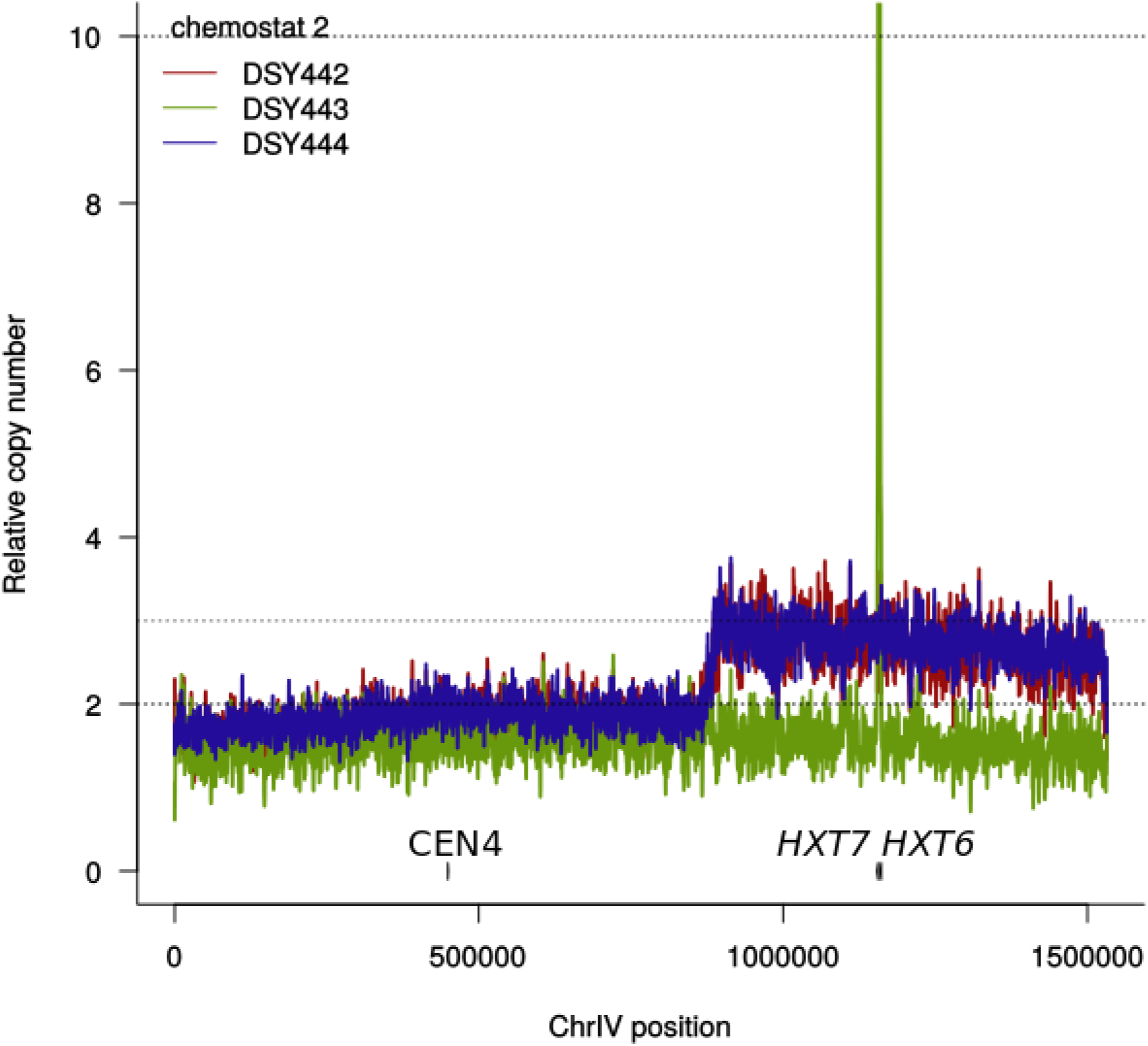
Relative coverage across chromosome IV of the wild-type colony phenotypes from chemostat 2. Two clones had a large partial duplication of the right arm of the chromosome (chrIV CNV) and the third had an approximately 10-fold expansion of the high-affinity hexose transporters *HXT6* and *HXT7* (*HXT6/7* CNV). Lines correspond to running medians of 801bp windows.

**Figure S6.**
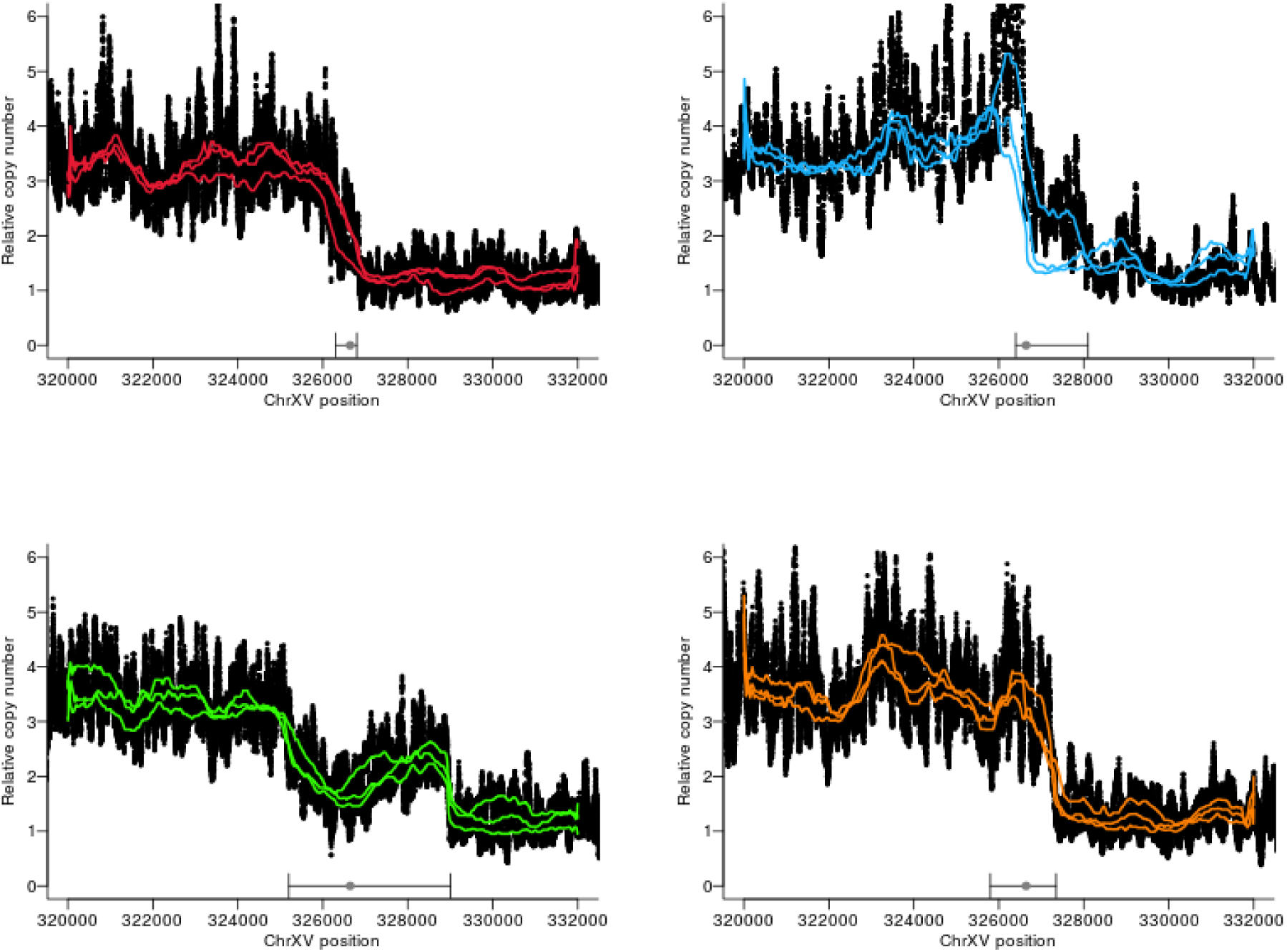
Relative coverage across chromosome XV of the small colony size phenotype strains from each chemostat. The circle and lines on the x-axis indicate the centromere location and the rearrangement boundaries (read pairs with same direction). Dots correspond to the relative coverage, while continuous lines to running medians of 801bp windows.

**Figure S7.**
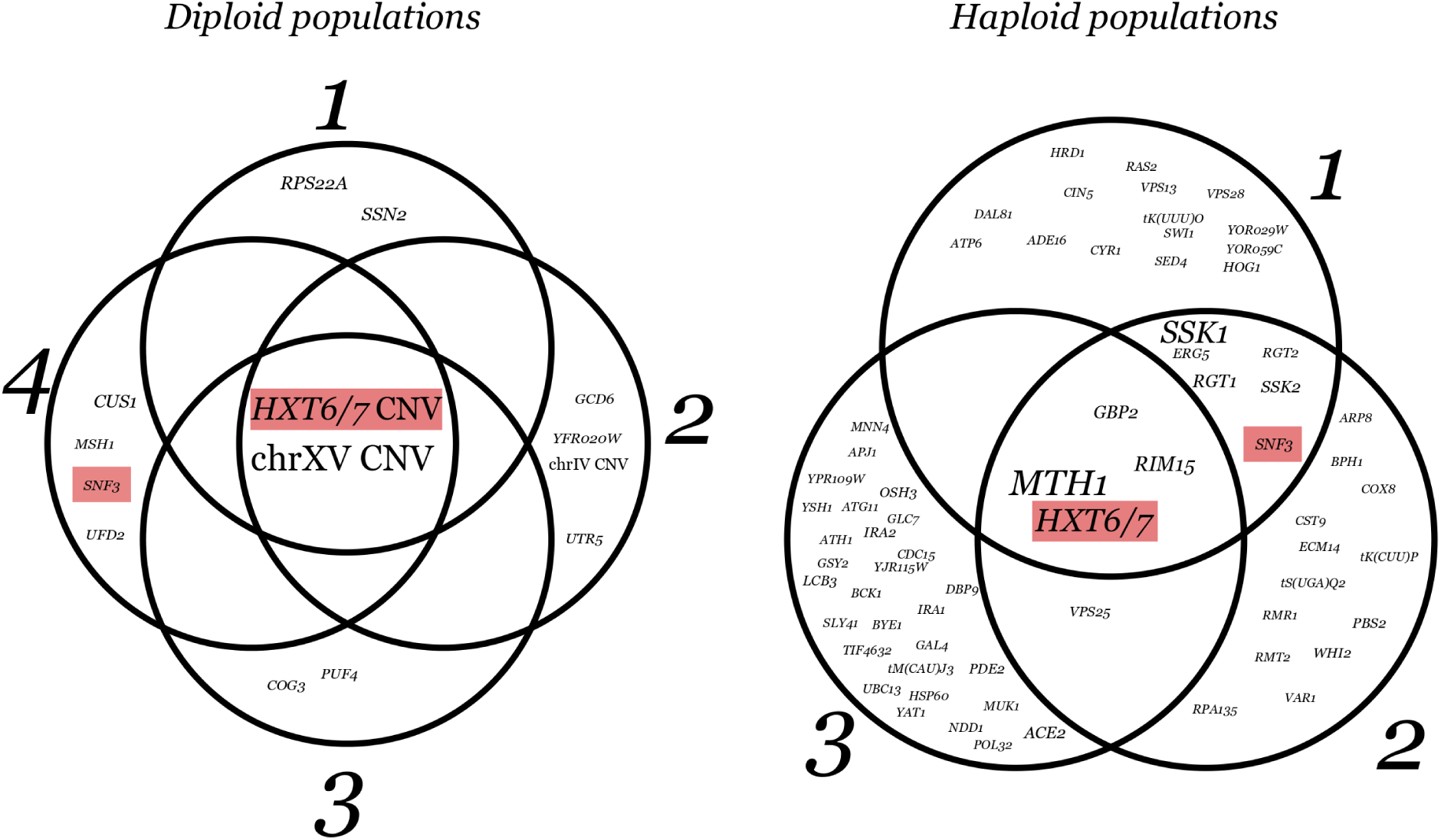
In both haploid and diploid replicate evolutionary experiments, a small number of genes are repeatedly targeted. Each circle contains the mutations found in an evolved haploid or diploid population. The font size of genes in the Venn diagram for haploids is proportional to the number of mutations in the corresponding gene identified in (Kvitek and Sherlock 2013). The font size of mutations in the Venn diagram for diploids is proportional to the number of strains in which the mutation was found (this study). The overlap between haploid and diploid targets is generally very low with the exception of the *HXT6/7* CNV. The only other shared gene targeted in both haploids and diploids is *SNF3* shown in red. The *HXT6/7* CNV frequency was not estimated in (Kvitek and Sherlock 2013), however from (Kao and Sherlock 2008) we know that it is very frequent and probably found in most populations and thus we included it in the figure with an arbitrary large font size as present in all three populations.

**Figure S8.**
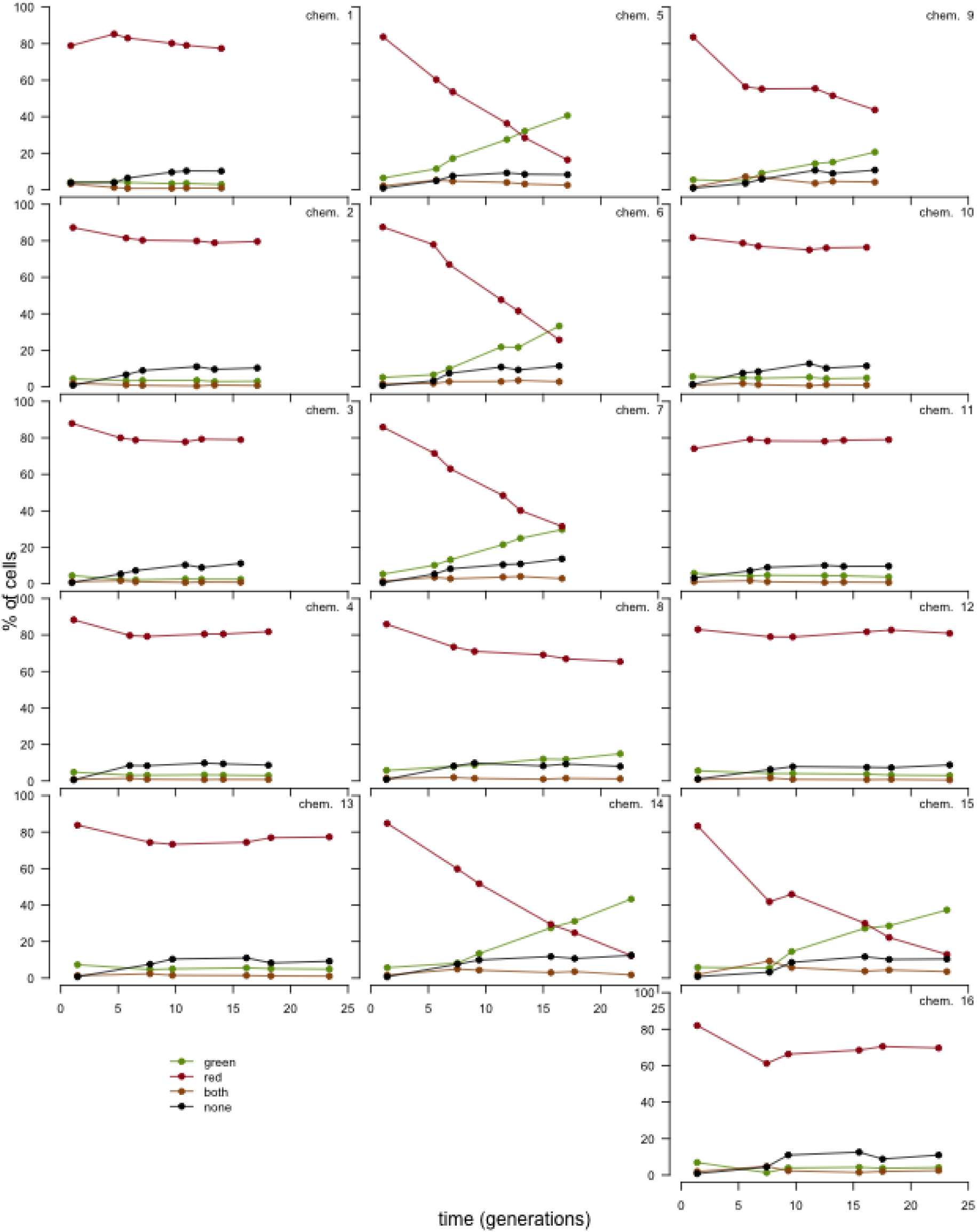
Frequency changes through time of the competing strains. Each plot corresponds to one chemostat. Each row of three plots consists of one biological replicate (three pairwise competitions). The first plot in each row corresponds to the ancestral genotype (-/-), the central to the heterozygote genotype (- / *HXT6/7* CNV) and the right to the homozygote (*HXT6/7* CNV / *HXT6/7* CNV) competing in each case against the red reference strain. For every time-point we FACS sorted 20,000 cells and each single cell measurement was classified as either carrying the green or the red fluorophore, or both or none. The double fluorescence (8.67% ± 4.44 mean and standard deviation across all measurements) is due to clumps of cells and measurement errors and the absence of fluorescence (2.23% ± 1.59) is either due to measurement error, dead cells or other detritus or stochastic lack of expression. Similar levels of double and no fluoresence were seen in control populations of pure cultures. The inocula (not included in the graph) had an average of 9.71 red to green ratio (± 2.33 standard deviation).

**Figure S9.**
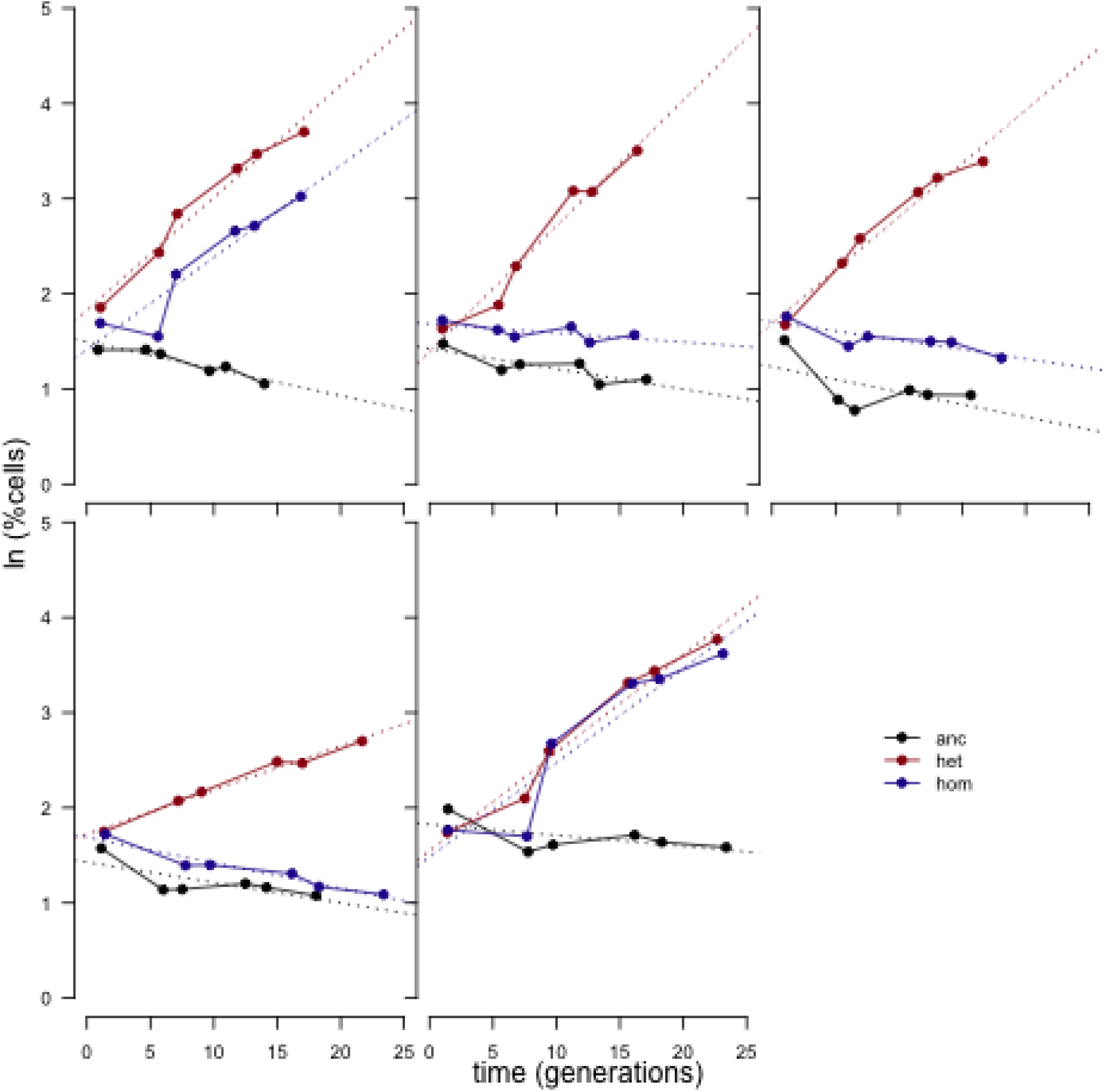
Relative selection coefficients of the ancestral homozygote genotype (-/-), the heterozygote (- / *HXT6/7* CNV) and the homozygote (*HXT6/7* CNV / *HXT6/7*). Each plot corresponds to one biological replicate (three pairwise competitions with the same reference). In four out of five replicates the heterozygote increases much faster than the homozygote. The dashed lines show the slope of the to linear regression.

